# AMF primes immune genes against *Puccinia hordei* (Brown rust) in *Hordeum vulgare* but does not reduce pathogen burden

**DOI:** 10.1101/2025.09.16.676302

**Authors:** Claire Moulton-Brown, Karolina Brzezinska, Beatriz Orosa-Puente, Thorunn Helgason

**Affiliations:** Institute of Ecology & Evolution, School of Biological Sciences, University of Edinburgh, Edinburgh, EH9 3BF, UK; Institute of Molecular Plant Sciences, School of Biological Sciences, University of Edinburgh, Edinburgh, EH9 3BF, UK

## Abstract

Barley (*Hordeum vulgare*) is susceptible to *Puccinia hordei* (leaf rust), a biotrophic foliar pathogen contributing to global yield losses. With rising food demand and increasing disease pressure, sustainable crop protection strategies are urgently needed to support UN Sustainable Development Goal 2: Zero Hunger. Arbuscular mycorrhizal (AM) fungi, including *Rhizophagus irregularis*, form symbioses with barley roots and can modulate host immunity through mycorrhiza-induced resistance (MIR). Here, we tested whether *R. irregularis* colonisation alters barley growth and defence responses during *P. hordei* infection.

AM fungal colonisation did not significantly reduce disease severity or mitigate pathogen-associated biomass loss at a single post-infection time point. However, co-infected plants showed enhanced expression of defence genes (*PR1*, *PR2*, *PR3* and *WRKY28*), which remained low in plants colonised by AM fungi alone, consistent with immune priming. RNA sequencing revealed AM fungal-associated reprogramming of the leaf transcriptome, including enrichment of defence, metabolism and ubiquitination-related processes.

These results indicate that *R. irregularis* reshapes barley immune regulatory networks at transcriptional and post-translational levels. Although these molecular changes did not translate into measurable phenotypic protection within the short experimental timeframe, they highlight the complexity and context dependence of MIR in cereal–rust interactions.

## Introduction

Food security faces mounting pressure as global demand is projected to rise by 35–56 % by 2050 (van Dijk *et al*. 2021). Crops are fundamental to global food security, yet pathogens already reduce yields by 20–40 % (Ristaino *et al*. 2021). Barley (*Hordeum vulgare*), the world’s fourth most cultivated cereal, is highly vulnerable to leaf rust (*Puccinia hordei),* which can reduce yield by up to 62 % in susceptible cultivars (Cotterill *et al*. 1992). Epidemics are driven by mild winters, wet springs and warm summers; conditions that are predicted to become more frequent with climate change (West *et al*. 2012). Current management relies on partially resistant cultivars, removal of volunteer hosts and fungicide application. However, these strategies are not fully effective as they are constrained by limited genetic resistance in UK barley (maximum Agriculture and Horticulture Development Board (AHDB) ratings of 6.2 for winter and 4.2 for spring varieties; AHDB 2025) and present sustainability concerns. Therefore, new approaches are needed to enhance crop resilience and reduce reliance on chemical inputs, in line with UN Sustainable Development Goal 2 (Zero Hunger and promotion of sustainable agriculture).

Arbuscular mycorrhizal (AM) symbiotic fungi represent an important strategy for promoting sustainable agriculture and enhancing food security. AM fungi (subphylum Glomeromycotina) colonise the roots of most vascular plants, including cereals (Wang and Qiu 2006). They facilitate nutrient and water uptake in exchange for sugars/lipids, leading to improved crop growth and yield, particularly under resource-limited conditions. Meta-analyses show that AM fungal inoculation increases cereal grain yields by up to 23 % under rainfed systems (Wu *et al*. 2022). Beyond nutrition, AM fungi also enhance tolerance to abiotic stressors such as drought and contribute to soil health via improved structure and microbial activity (Diagne *et al*. 2020).

At the molecular level, symbiosis with AM fungi has been shown to trigger extensive transcriptional reprogramming in host plants, including the activation of genes involved in nutrient transport, hormone signalling, and stress adaptation (Gao *et al*. 2023; Gong *et al*. 2023). Crucially, AM fungi can reduce susceptibility to pests and pathogens by modulating host immunity in a phenomenon known as mycorrhiza-induced resistance (MIR) (Farhaoui *et al*. 2025). The use of AM fungal symbioses is proposed as a sustainable strategy to improve cereal crop productivity while reducing dependence on synthetic fertilisers and pesticides (Bender, Wagg and van der Heijden 2016; Umer *et al*. 2025). To do this effectively, we must understand the mechanisms that underpin these processes, so that either management strategies that enhance AM function (Austen *et al*. 2022), or appropriate inoculants can be identified that genuinely enhance productivity (Elliot *et al*. 2020).

Mycorrhiza-induced resistance (MIR) provides systemic protection against a broad range of pathogens and shares similarities with both pathogen-triggered systemic acquired resistance and rhizobacteria-induced systemic resistance (Cameron *et al*. 2013). Rather than triggering constitutive defence activation, MIR predominantly functions through immune priming, whereby defence responses are sensitised and more rapidly or strongly deployed upon pathogen challenge, minimising the fitness costs associated with sustained immune activation (Jung *et al*. 2012).

AM fungal colonisation remodels host immune signalling networks, enabling modified defence responsiveness at sites distinct from the roots. Hormonal crosstalk plays a central role in this process. Early models suggested that AM fungi suppress salicylic acid (SA)-dependent defences while enhancing jasmonic acid (JA)-mediated responses, conferring resistance to necrotrophs and herbivores (Cameron *et al*. 2013). However, more recent findings indicate that MIR can amplify both SA- and JA-responsive genes upon pathogen challenge, showing a more integrated modulation of multiple defence pathways (Fujita *et al*. 2022).

Importantly, MIR is systemic in nature, with AM fungal colonisation inducing transcriptional reprogramming not only in roots but also in distal tissues such as leaves, where altered defence signalling and priming responses to foliar pathogens are ultimately expressed (Pozo *et al*. 2002). The MIR phenotype has been reported across diverse plant taxa. In tomatoes (*Solanum lycopersicum*), *Funneliformis mosseae* colonisation enhanced resistance to *Alternaria* blight through JA-dependent signalling, and protection was lost in JA-pathway mutants (Song *et al*. 2015). In wheat (*Triticum aestivum*), AM fungal colonisation with *F. mosseae* reduced susceptibility to powdery mildew (*Blumeria graminis*) via enhanced accumulation of defence metabolites and upregulation of pathogenesis related (PR) and phenylpropanoid biosynthetic genes (Mustafa *et al*. 2017). Similarly, *F. mosseae* was found to prime wheat against *Zymoseptoria tritici* (Allario *et al*. 2025), and *Rhizophagus irregularis* improves rice (*Oryza sativa*) resistance to *Magnaporthe oryzae,* which in turn increased grain yield in field settings (Martín-Cardoso *et al*. 2025). These cases demonstrate the relevance of MIR to both monocot and dicot systems and highlight its potential to enhance disease resistance and yield concurrently, a central goal of sustainable crop protection.

Mechanistically, MIR involves transcriptional and post-transcriptional reprogramming of immune signalling networks. Mycorrhizal colonisation alters the expression of transcription factors, including WRKY proteins, which regulate both basal and inducible defences. In apple (*Malus domestica*), AM fungal-induced *WRKY40* enhanced resistance to *Fusarium* by activating β*-1,3-glucanase* expression (Wang *et al*. 2022). Priming also enhanced physical defences, such as callose deposition, which restricts pathogen entry. Blocking callose synthesis abolishes MIR protection in tomato (Sanmartín *et al*. 2021), indicating its functional importance.

Beyond transcriptional changes, MIR includes modulation of the ubiquitin-proteasome system (UPS), a major post-translational regulator of immunity. E3 ubiquitin ligases, which confer substrate specificity in UPS, target immune regulators for proteasomal degradation, balancing activation and repression of defences (de Vega, Newton and Sadanandom 2018). Several *PUB* and *Arabidopsis Toxicos en Levadura* (*ATL*) family ligases are upregulated in mycorrhizal roots and shoots (Wu *et al*. 2024), suggesting that AM fungi may fine-tune immune signalling by modifying proteostasis. This regulation may allow plants to remain immunologically alert without incurring the growth penalties of constitutive defence.

The magnitude and outcome of MIR can be highly context dependent. Factors including plant nutrient status, AM fungal genotype, pathogen lifestyle, and environmental conditions can influence whether mycorrhiza-induced changes translate into effective resistance (Dejana *et al*. 2022; Weinberger *et al*. 2025). Understanding how these variables interact in cereal systems is vital if MIR is to be harnessed as a reliable strategy for sustainable crop protection. Despite compelling evidence from other pathosystems, the effectiveness and molecular basis of MIR in barley, particularly against biotrophic foliar pathogens like *P. hordei*, remains unstudied. Evidence from cereal-pathogen-symbiont tripartite systems is needed to validate MIR as a practical tool for disease management. In this study, we hypothesised that colonisation of barley by *Rhizophagus irregularis* (a widely distributed AM fungus) primes the barley immune system, which could attenuate *P. hordei* infection or disease development.

First, we quantified barley growth traits and rust disease severity to assess whether arbuscular mycorrhizal fungal colonisation influences host fitness or pathogen susceptibility. These measurements directly test the hypothesis that AM fungi enhance tolerance or resistance to *Puccinia hordei* without incurring growth penalties, a hallmark of priming-based immune responses. Second, we examined basal and pathogen-induced expression of key defence-associated gene families, including pathogenesis-related (PR) proteins and WRKY transcription factors. These gene groups were selected as central regulators of plant immunity. Alterations in the basal expression of plant immune markers are indicative of a primed state, whereas changes in induction kinetics or amplitude upon pathogen challenge reflect enhanced defence activation. Third, we conducted transcriptomic analyses of leaf tissue to capture systemic reprogramming of defence-related pathways in response to AM fungal colonisation. This enabled assessment of broader immune signalling networks, including transcriptional regulation and ubiquitin-mediated protein turnover, which are increasingly recognised as key components of immune priming and defence fine-tuning.

By integrating physiological phenotyping with targeted and global transcriptional analyses, this study aims to assess whether *R. irregularis* can induce mycorrhiza-induced resistance (MIR) in the barley-*P. hordei* pathosystem and to characterise its potential mechanistic basis. Given emerging evidence that MIR outcomes and underlying regulatory networks can vary depending on host species, microbial symbionts, and pathogen lifestyles, dissecting MIR in this economically relevant cereal-rust interaction is critical for distinguishing conserved immune features from system-specific regulatory mechanisms. Such comparative resolution is essential for assessing the translational potential of microbiome-based resistance strategies across diverse crop-pathogen combinations.

## Materials and Methods

### Experimental design

A 2 × 2 factorial experiment was conducted to assess the impact of AM fungal colonisation and foliar pathogen infection on barley. Treatments comprised: (i) **C**ontrol (no AM fungi, no pathogen), (ii) **A**M fungi only (*Rhizophagus irregularis*), (iii) ***P****uccinia hordei* only, and (iv) both AM fungi and *P. hordei* (**AP**). Each treatment group included six biological replicates, with a subset of three replicates used for transcriptomic analysis and ubiquitination quantification. Barley was grown for four weeks, then inoculated with *P. hordei* spores. The experiment was terminated one week post-inoculation (Figure 1), a key timepoint for the detection of differences in infection establishment and defence gene induction that are most informative for elucidating MIR mechanisms (Arntzen and Parlevliet 1986).

**Figure 1.**
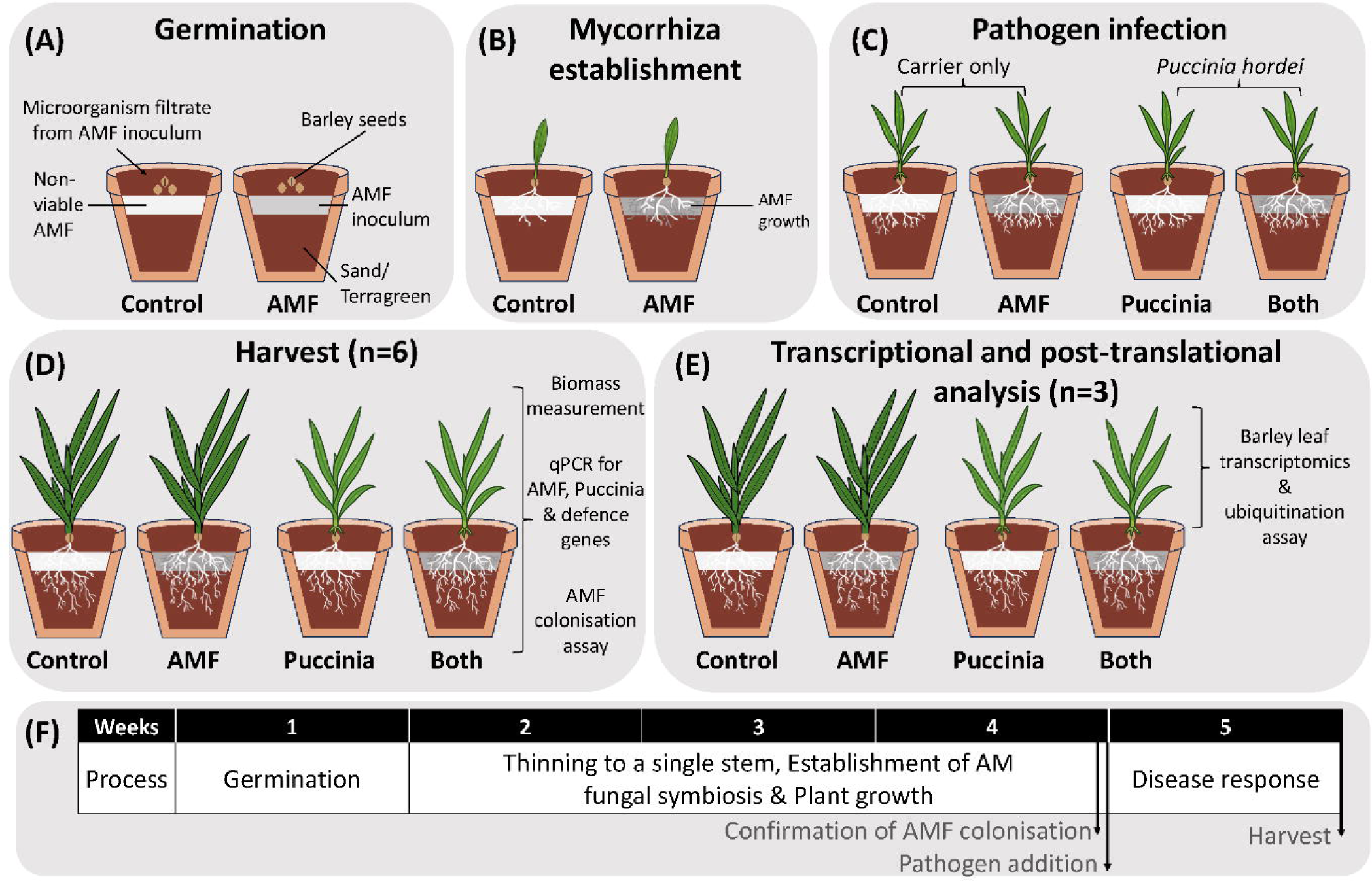
Overview of the experimental setup showing (A) the treatment setup, (B) the AMF colonisation phase when pots were thinned to a single stem, (C) *Puccinia hordei* inoculation, (D) the final harvest, including measurements of plant growth, AMF colonisation and qPCR-based quantification of AMF, Puccinia and defence gene expression (n = 6), (E) transcriptional and post-translational analyses performed on a subset of plants (n = 3) with barley leaf transcriptomics and ubiquitination assay and (F) the experimental timeline. Figure adapted from Weinberger et al. (2025).

### Plant growth conditions and AM fungal inoculation

Seeds of *Hordeum vulgare* cv. Bowman, a rust-susceptible cultivar, were surface sterilised by vortexing in 30 % (v/v) bleach supplemented with 0.1 % (v/v) Triton X-100 using three sequential incubation steps (15 s vortex followed by 6 min at room temperature, repeated twice, followed by a final 20 s vortex). Seeds were then washed three times with sterile deionised water and sown in a 1:1 (v/v) mixture of autoclaved coarse silica sand and Terragreen® clay (Oil-Dri, UK). Each 1 L pot was supplemented with 0.5 g L⁻¹ bonemeal and watered twice weekly with phosphate-free Hoagland’s solution to promote mycorrhization.

Plants were grown in a controlled-environment chamber (MLR-352H, PHC, Japan; 20°C day / 17°C night; 70 % relative humidity) under a 16 h light/8 h dark photoperiod, with fluorescent lighting (FL40SSENW37) providing 150 µmol m⁻² s⁻¹.

AM fungi treated plants received commercial *Rhizophagus irregularis* inoculum (PlantWorks, UK) at 20 g L⁻¹ of growth substrate, a rate previously shown to ensure consistent and robust AM fungal colonisation in controlled experiments using sterilised media (Dejana *et al*. 2022; Weinberger *et al*. 2025). Controls received autoclaved inoculum and microbial filtrate to standardise non-AM fungal microbial communities.

### Pathogen inoculation

*Puccinia hordei* isolate BBR 20-003 was obtained from the National Institute of Agricultural Botany (NIAB). Urediniospores (∼1.3 × 10LJ spores plant⁻¹) were applied to leaves using talcum powder as an inert carrier (Roelfs, Singh and Saari 1992) at a spore: talcum ratio of 1:10 (w/w). Spores were mixed with talcum powder immediately prior to inoculation and dusted onto the adaxial leaf surface using a fine brush to ensure uniform distribution. Control plants received talcum powder only. To promote infection and prevent cross-infection, *P. hordei*-inoculated plants were maintained at >95 % relative humidity for 12 hours post-inoculation and transferred to a separate growth cabinet under identical growth conditions.

### Confirmation of AM fungal colonisation

AM fungal colonisation was verified by both microscopy and qPCR on root samples. For microscopy, roots were stained using the ink and vinegar method (Vierheilig *et al*. 1998). Roots were stored in 40 % ethanol at 4°C, then cleared with 10 % KOH at 70°C for 45 minutes. They were stained with 5 % Brilliant Black ink (Pelikan) in 5 % acetic acid, destained in 1 % acetic acid, and mounted on slides with 50 % (v/v) glycerol.

For AM fungal qPCR, DNA was extracted from roots using the DNeasy Plant Mini Kit (Qiagen). Each 10 µL qPCR reaction included Fast SYBR™ Green Master Mix (Thermo Fisher Scientific), 350 nM of each primer, and ∼10 ng of root DNA. AM fungi were quantified using the 18S rRNA gene with AM fungal-specific primers AMG1F and AM1 (Table S1; Helgason et al. 1998; Hewins, Carrino-Kyker and Burke 2015), normalised to barley Elongation Factor 1-α using primers HvEF1a-qF and HvEF1a-qR (Hua *et al*. 2015). Triplicate reactions were run on a 7500 Fast Real-Time PCR System. The program comprised an initial denaturation at 95°C for 20 s, followed by 40 cycles at 95°C for 10 s and 60°C for 30 s, then a melt curve from 60-95°C. Data were analysed with Applied Biosystems 7500 Software (v2.3) using the Pfaffl method for relative quantification.

### Rust infection quantification

Pathogen biomass in leaves was quantified by RT-qPCR (see Gene expression analysis below for conditions). Leaf RNA was extracted and converted to cDNA (see below). Primers targeting the *P. hordei* elongation factor 2 gene (Table S1) were used to measure pathogen cDNA relative to barley cDNA (using a barley endogenous actin gene as reference). This provided a relative infection index for each plant.

### Barley resource allocation measurements

Root and shoot dry weights were recorded at harvest. Plants were divided into above- and below-ground fractions and dried at 60°C until a constant weight was reached. Total biomass and root:shoot ratio were calculated.

### RNA extraction and cDNA synthesis

Total RNA was extracted from barley leaf tissue using a phenol-chloroform-isoamyl alcohol method with LiCl precipitation, adapted from established protocols (Verwoerd, Dekker and Hoekema 1989). Briefly, frozen plant material was ground under liquid nitrogen, extracted with warmed phenol : chloroform : isoamyl alcohol (25 : 24 : 1), and subjected to sequential organic washes. The aqueous phase was precipitated with 1/3 volume of 8 M LiCl overnight at 4°C, and the resulting pellet was washed with 70 % ethanol. A secondary precipitation with sodium acetate and 96 % ethanol was performed to ensure purity, and RNA pellets were finally resuspended in RNase-free water. cDNA was synthesised using the SuperScript™ III Reverse Transcriptase (Invitrogen) according to the manufacturer’s instructions.

### Gene expression analysis

Quantitative RT-PCR was used to assess the expression of immune marker genes, *PR1*, *PR2*, *PR3*, and *WRKY28*. Reactions used SYBR™ Green Universal Master Mix (Applied Biosystems, Thermo Fisher Scientific). Reactions were conducted in technical triplicates, each containing 2 µL of cDNA and 200 nM gene-specific primers (Table S1). The reactions were run on a QuantStudio 5 system (Applied Biosystems, Thermo Fisher Scientific). The cycling conditions were: 50°C for 2 minutes and 95°C for 10 minutes; 40 cycles of 95°C for 15 seconds, 60°C for 1 minute; and a melt curve at 95°C for 15 seconds, 60°C for 1 minute, and 95°C for 15 seconds. Relative expression was calculated by the ΔCt method and normalised to the barley housekeeping gene (Actin).

### RNA sequencing and transcriptomics

RNA sequencing was conducted on a subset of samples (n = 3 per treatment). RNA sequencing was performed by Novogene Company Limited using the Illumina NovaSeq Platform (150 bp paired end). Reads were quality assessed using FastQC v0.11.9 (Andrews 2010). Adapter sequences were trimmed using Trimmomatic v0.39 (Bolger, Lohse and Usadel 2014). After adapter and quality trimming, we obtained on average 43.69, 42.83, 32.90, and 44.11 Mb clean reads from the Control, AM fungi, *P. hordei*, and AM fungi plus *P. hordei* treatments, respectively (Table S2), with a similar GC content across treatments (53.34 %). RNA-sequencing reads were aligned and annotated against the barley genome (MorexV3) using HISAT2 v2.1.0 (Kim *et al*. 2019).

### Ubiquitination assay

Given the enrichment of ubiquitin-related genes in AM fungal-treated plants, global protein ubiquitination was assessed by Western blot. Total protein was extracted from frozen barley leaves using a RIPA-like buffer adapted from established plant protocols (Chico *et al*. 2020). Briefly, tissue was ground under liquid nitrogen and homogenised in extraction buffer (20 mM Tris pH 7.5, 50 mM NaCl, 1 mM EDTA, 0.5 % NP40, 0.5 % SDS, 0.5 % sodium deoxycholate) supplemented with protease inhibitors, 50 µM MG132, and 10 mM N-ethylmaleimide (NEM). Extracts were vortexed, incubated on ice for 10 min, and centrifuged at 10,000 rpm for 15 min at 4°C. The supernatant was collected for downstream analysis.

Protein extracts were heated in SDS sample buffer with 50 mM DTT at 80°C for 10 minutes and resolved by SDS-PAGE. Proteins were transferred to a nitrocellulose membrane via wet transfer at 20 V overnight. Transfer efficacy was verified with Ponceau staining. Membranes were blocked with 5 % (w/v) skim milk powder (Millipore) in PBST for 2 hours, then incubated with primary antibodies (Poly-ubiquitin P4D1 at 1:2500, Santa Cruz Biotechnology; Actin at 1:5000, Agrisera; actin was used as a loading control for normalisation) for 1.5 hours, followed by three 5-minute washes in PBST. Secondary antibody, anti-mouse 1:5000 (Cell Signaling Technology), incubation lasted 1 hour, followed by three additional washes. Detection was performed using a chemiluminescent substrate (SuperSignal™ West Pico PLUS, Thermo Fisher Scientific) and visualised with LI-COR® Odyssey Fc Imager with band intensity quantified using Image Studio 6.1 (LICORbio).

### Statistical analysis

Statistical analyses were performed using R v4.5.1 (R Core Team 2025) implemented in RStudio version 2025.09.2 (RStudio Team 2025). Data processing, reshaping, and visualisation were performed using tidyverse v2.0.0 (Wickham *et al*. 2019), primarily dplyr v1.1.4, tidyr v1.3.1, and ggplot2 v4.0.1, with multi-panel figures assembled using cowplot (v1.2.0).

Physiological traits (root dry mass, shoot dry mass, and root:shoot ratio) and quantitative PCR (qPCR) data were analysed using two-way analysis of variance (ANOVA) to test for the main effects of arbuscular mycorrhizal fungal (AM fungal) inoculation (AM-, AM+) and *Puccinia hordei* infection (Puccinia-, Puccinia+), and their interaction. ANOVA models were fitted using the base R function aov() from the stats package, with F-tests and associated p-values obtained using anova(). qPCR data were log2-transformed prior to statistical analysis. Post-hoc comparisons among factorial treatment combinations were conducted using estimated marginal means calculated with the emmeans() function from the emmeans package v2.0.0 (Russell V. Lenth and Julia Piaskowski 2025). Pairwise comparisons were adjusted for multiple testing using the Sidak method, and statistically distinguishable groups were summarised using compact letter displays generated with the cld() function from the multcomp package v1.4-29 (Hothorn, Bretz and Westfall 2008). Exact p-values are reported in figures, with significance indicated by asterisks (* p < 0.05, ** p < 0.01, *** p < 0.001).

RNA gene count data were quantified and standardised using HTseq2 v2.0.5 (Putri *et al*. 2022). Differential analysis was performed using edgeR v3.42.4 (Chen *et al*. 2025) by filtering out CPM < 1 and normalising using a TMM method (Robinson and Oshlack 2010). Differentially expressed genes (DEGs) were defined as: FDR < 0.05, log_2_FC > 1, logCPM > 2. Gene Ontology (GO) enrichment analysis of DEGs was done using g:Profiler (Raudvere *et al*. 2019) with FDR correction. Hierarchical clustering (Ward’s method) of DEGs was employed to identify groups of genes with similar expression patterns across the four conditions.

Ubiquitination data were normalised to actin and expressed as fold change relative to the control treatment. Statistical analyses of ubiquitination data were performed on log₂-transformed fold-change values using the same two-way ANOVA framework described above.

Assumptions of normality and homoscedasticity were assessed by visual inspection of ANOVA residuals (residuals versus fitted values and normal Q-Q plots). Where necessary, data were transformed prior to analysis to improve adherence to model assumptions and stabilise variances. Independence of observations was ensured by the experimental design, with each plant treated as an independent experimental unit.

## Results

### AM fungi alone did not reduce rust infection or improve biomass under pathogen challenge

Barley plants were inoculated with *Rhizophagus irregularis*, and root colonisation was confirmed 28 days post-inoculation. Microscopy revealed abundant arbuscules and hyphae in the inoculated roots, whereas no mycorrhizal structures were observed in non-inoculated control samples. qPCR further validated successful AM fungal colonisation in inoculated plants (Figure 2 A-B). Low-level amplification detected in control samples is consistent with previously reported non-specific amplification by the AM1 primer in the absence of AM fungal templates (Lee, Lee and Young 2008). AM fungal presence did not significantly affect rust severity one week after *P. hordei* infection; *P. hordei*-specific marker gene (PhEF2) showed only a slight, non-significant increase in AM fungi-colonised plants (Figure 2 C).

**Figure 2.**
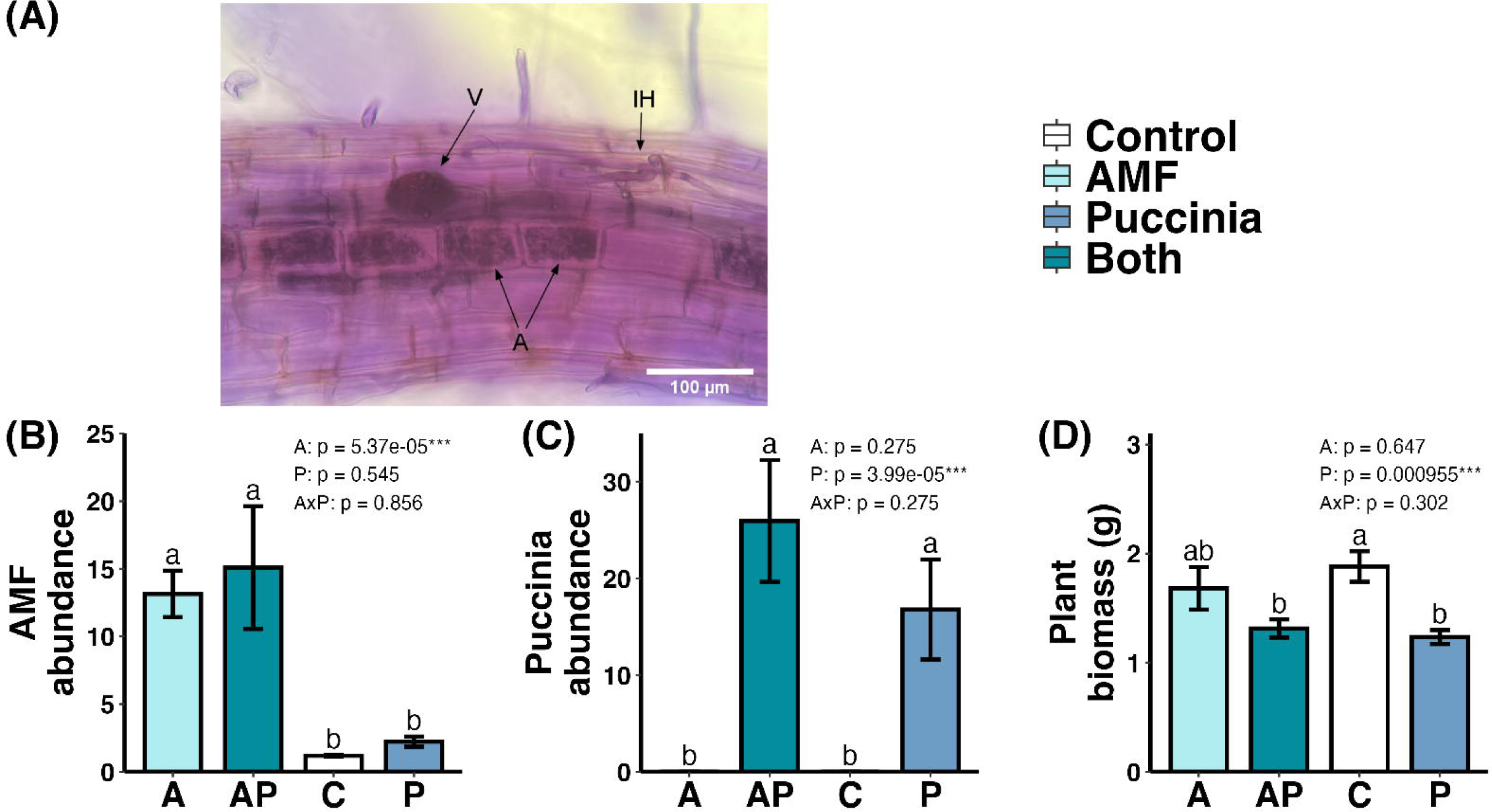
Endpoint measurements of AMF colonisation, *Puccinia* infection, and plant biomass after one week of *Puccinia hordei* infection. (A) Photograph of barley roots stained to show AMF colonisation, highlighting vesicles – V, intraradical hyphae – IH, and arbuscules – A. (B) Relative AMF DNA in barley roots, quantified as AMF copy number normalised to *HvEMF1*α, five weeks post-inoculation. (C) Relative abundance of *Puccinia* in barley leaves, quantified as *PhEF2* copy number normalized to *HvACT*, one-week post-infection. (D) Total dry biomass of shoots and roots after five weeks of growth. Bars sharing different letters indicate significant differences (p < 0.05). C, control; A, AMF addition; P, *Puccinia* addition; AP, AMF plus *Puccinia* addition (mean ± SE, n = 6).

*Puccinia hordei* infection resulted in a substantial reduction in plant biomass, with all infected plants exhibiting approximately 20% lower total dry weight than uninfected controls, regardless of AM fungal colonisation (p = 0.000955; Figure 2 D). No significant AM fungi × pathogen interaction was detected, and the root:shoot ratio remained unchanged (Figure S1), indicating a proportional reduction in overall growth rather than altered biomass allocation. In contrast, AM fungal colonisation had no detectable effect on barley growth (Figure 2 D and S1), indicating that AM fungi did not impose a growth penalty under these experimental conditions.

Overall, *P. hordei* caused substantial biomass losses regardless of AM fungal presence. We next assessed whether AM fungal colonisation was associated with changes in molecular immune responses that were not evident in infection or growth metrics in this timeframe.

### AM fungi primed the expression of barley defence genes during *P. hordei* infection

Although AM fungi did not reduce rust severity in this experiment, we tested whether AM colonisation altered defence gene expression patterns during infection. Transcript levels of SA-responsive PR genes (*PR1*, *PR2*, *PR3*) and the transcription factor *WRKY28* were measured. In the absence of *P. hordei*, *PR1*, *PR2* and *PR3* transcript levels remained low in AM-colonised leaves and did not differ from non-colonised controls (Figure 3 A-C).

**Figure 3.**
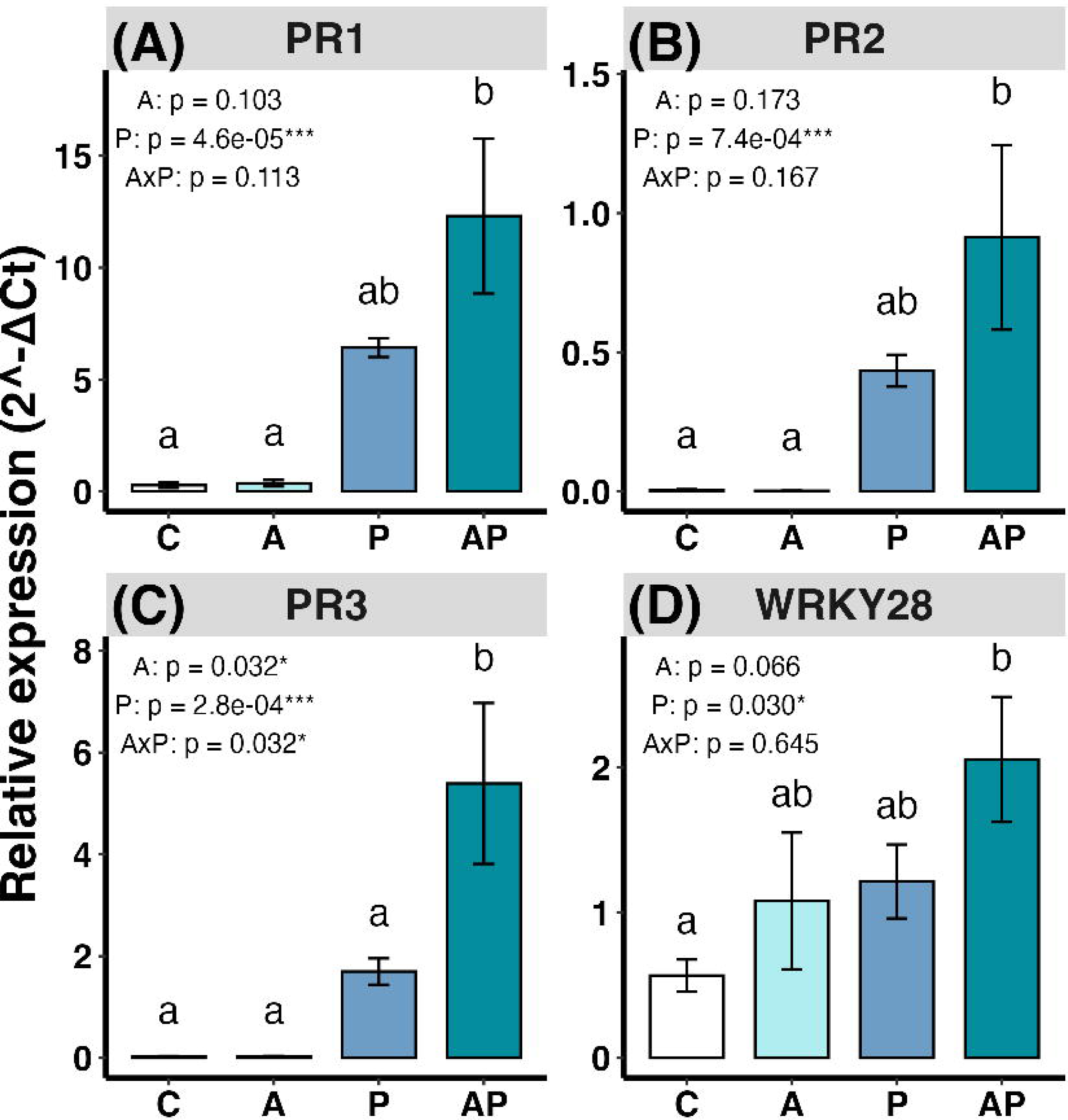
Relative expression of barley immune marker genes (A) *PR1*, (B) *PR2*, (C) *PR3*, (D) *WRKY28*. Gene expression is relative to barley actin (HvACT); Bars sharing different letters indicate significant differences (P < 0.05). C, control; A, AMF addition; P, *Puccinia* addition; AP, AMF plus *Puccinia* addition (mean ± SE, n = 6).

Following *P. hordei* infection, *PR1*, *PR2* and *PR3* were induced, with higher transcript levels in AM-colonised infected plants than in infected plants without AM fungi (Figure 3 A-C). *WRKY28* showed a similar pattern, with low expression in uninfected plants and increased expression following infection, including higher transcript levels in AM fungi-colonised infected plants (Figure 3 D). This pattern is consistent with enhanced activation of higher-order immune regulators.

Together, these results show that priming occurred at the transcriptional level. AM fungi alone did not trigger immune gene expression in the absence of a pathogen but enhanced the amplitude of the transcriptional response upon infection. The lack of immediate protection despite strong PR induction suggests that *P. hordei* continues to suppress or evade host defences, or that timing and physiological constraints limit the effectiveness of the primed response.

### AM fungi reprogrammed the barley leaf transcriptome

For a comprehensive view of the transcriptomic changes induced by AM fungi, we performed RNA sequencing of barley leaves. AM fungal colonisation altered the expression in healthy leaves of 526 genes (344 upregulated, 182 downregulated; Figure 4 A; Figure S2). A Gene Ontology (GO) enrichment analysis revealed that upregulated genes were enriched for defence, stress responses, and protein modification (Figure 4 B). Additionally, upregulated transcripts included sugar transporters and amino acid biosynthesis genes, whereas downregulated transcripts included those annotated for growth, photosynthesis and gibberellin signalling (Table S3). This suggests AM fungi enhance the host’s capacity to regulate protein turnover. Induction of sugar transporters and amino acid synthesis genes likely reflects increased nutrient exchange, while downregulated transcripts related to growth, photosynthesis, and gibberellin signalling indicate a moderated growth programme. These shifts did not affect overall biomass (Figure 2 D).

**Figure 4.**
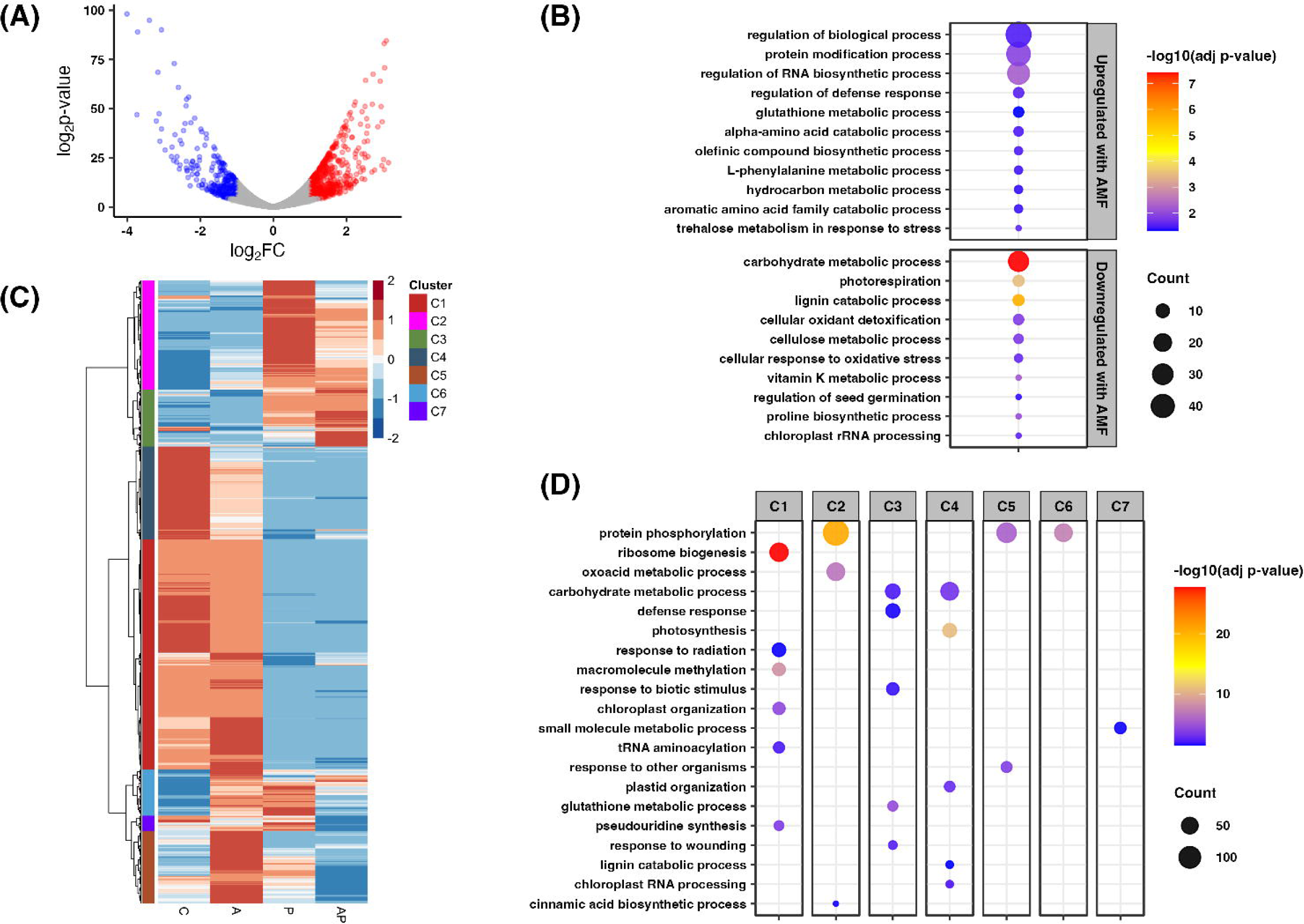
Transcriptional responses to AMF colonisation and *Puccinia* infection in barley leaves (n = 3). (A) Differentially expressed genes between control and AMF-colonised plants (Control vs AMF; log_2_FC > 1, FDR < 0.05, logCPM > 2). (B) GO enrichment of AMF-responsive genes shown in (A), using g:Profiler with Benjamini-Hochberg correction (adjusted *P* < 0.05, term size > 2). (C) Heatmap of row-wise z-scored log_2_-transformed expression [log_2_(x + 1)] for genes differentially expressed across treatments (C, control; A, AMF addition; P, Puccinia addition; AP, AMF and Puccinia), clustered into seven co-expression groups. (D) GO enrichment of gene clusters shown in (C), using g:Profiler with g:SCS correction (adjusted *P* < 0.05).

*P. hordei* infection in non-mycorrhizal plants triggered differential expression of 4,258 genes (P/C: 2,317 upregulated, 1,941 downregulated), indicating a robust transcriptional reprogramming in response to the pathogen. In comparison, AM fungal colonisation attenuated the overall magnitude of rust-induced gene expression changes, reducing the number of upregulated genes by ∼15 % and downregulated genes by ∼30 % (AP/A, 1,956 upregulated, 1,380 downregulated). Interestingly, during *P. hordei* infection, 231 genes were differentially expressed in AM-colonised plants compared with non-mycorrhizal plants (AP/P, 163 upregulated, 68 downregulated), suggesting that AM fungi still shape a distinct transcriptional response. Cluster analysis revealed that AM fungi amplified defence-related genes, including PRs and glutathione-associated transcripts (Cluster 3), while dampening broad kinase- and receptor-driven signalling cascades (Clusters 2 and 4; Figure 4 C-D). Genes induced by AM fungi alone but suppressed during co-infection (Clusters 5 and 6) may reflect negative feedback on pre-activated signalling modules. Together, these results suggest that AM fungi modulate transcriptional changes by limiting broad stress responses and enhancing defence-related gene expression.

### AM fungi altered defence-associated signalling and transcriptional responses

To further investigate AM fungal-mediated modulation of plant immune response, we examined key components of canonical immune signalling pathways. Transcriptional reprogramming is a hallmark of immune activation and our targeted qPCR analyses had already indicated that AM fungal colonisation influenced the expression of the immune-associated transcription factor *WRKY28* (Figure 3 D), which showed enhanced induction in AM fungi-colonised plants during *Puccinia* infection. This observation prompted a broader investigation of transcription factor dynamics at the transcriptome scale.

We therefore focused on the WRKY transcription factor family, which comprises major regulators of both biotic and abiotic stress responses (Javed and Gao 2023; Li *et al*. 2024) and has previously been shown to be regulated during AM colonisation (Gallou, Declerck and Cranenbrouck 2012). We found 74 annotated barley WRKY genes that were differentially expressed across treatments (Table S4). In healthy plants, AM fungi generally upregulated the expression of WRKY transcription factors, with 95 % of differentially expressed WRKY genes showing increased expression. In contrast, during *P. hordei* infection, AM fungi suppressed the majority of rust-induced WRKYs, downregulating 90 % of those that were otherwise induced by infection alone (Figure 5 A). Notable exceptions included *WRKY27*, *WRKY63*, and *WRKY1/38*, which were induced even more in co-infected plants. Most differentially expressed WRKYs belonged to Classes II and III, consistent with a role in biotic stress regulation.

**Figure 5.**
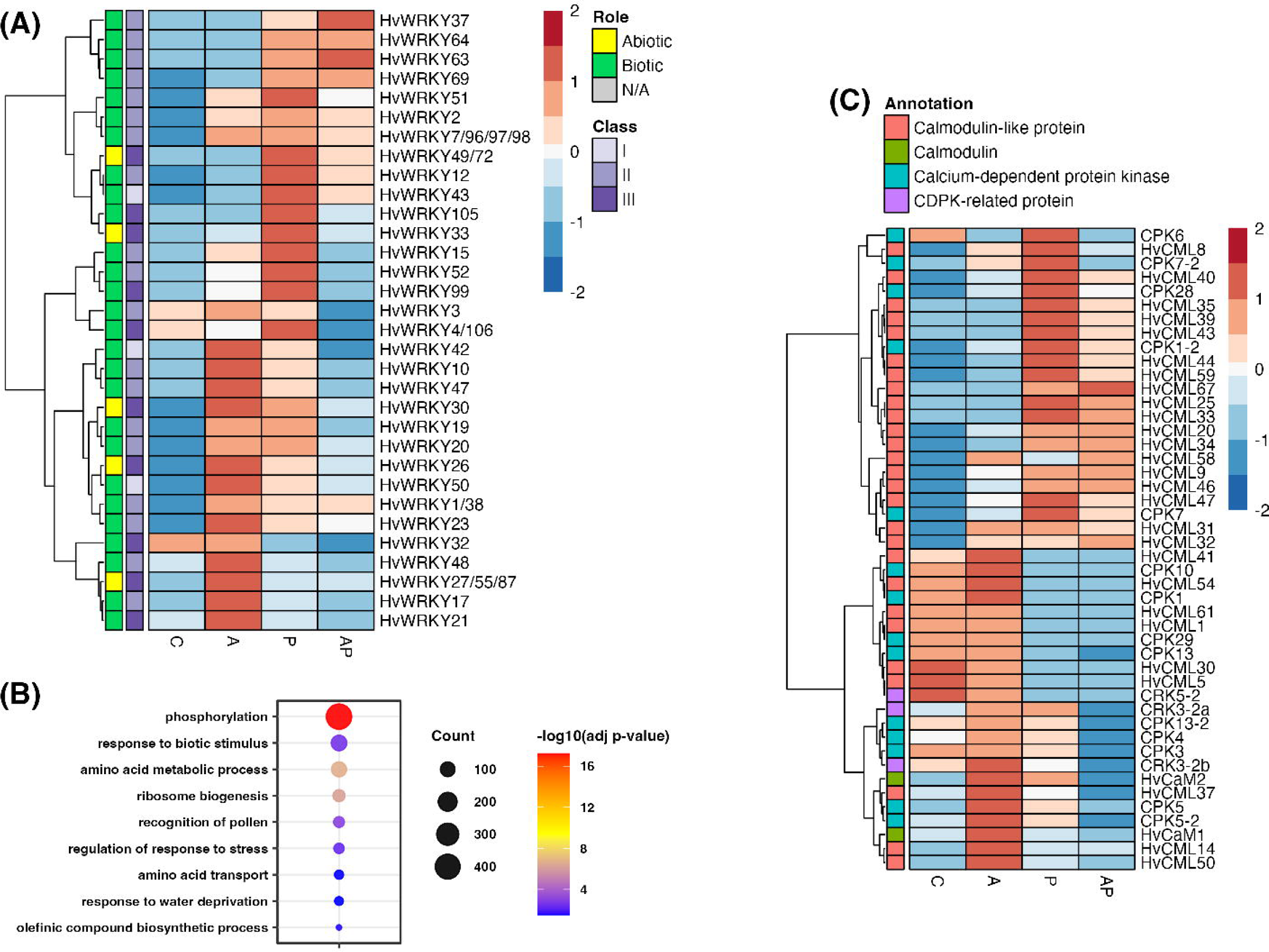
AMF modulation of WRKY transcription factor expression and calcium signalling in barley leaves (n = 3). (A) Row-scaled expression of differentially expressed WRKY genes across control (C), AMF (A), Puccinia (P), and combined (AP) treatments (log₂FC > 1, FDR < 0.05, logCPM > 2), annotated by WRKY class and stress-response role. (B) GO enrichment of differentially expressed genes containing predicted WRKY-binding motifs within 500 bp upstream promoter regions (FIMO, P < 0.001). (C) Row-scaled expression of differentially expressed calcium signalling genes across the same treatments (log₂FC > 1, FDR < 0.05, logCPM > 2). Calmodulin and calmodulin-like genes were annotated based on (Cai et al. 2022); calcium-dependent protein kinase (*CDPK*) and *CDPK*-like genes were annotated based on their closest Arabidopsis homologues.

To assess whether autoregulation and cross-regulation by WRKY transcription factors contribute to the observed transcriptional changes, we then scanned the 500 bp upstream promoter regions of all DEGs for W-box elements (TTGACY), the canonical WRKY binding motif. Approximately 80 % of DEGs contained at least one W-box and gene ontology enrichment analysis showed their involvement in phosphorylation, alongside terms related to response to biotic stimulus, stress, and olefinic compound biosynthesis (Figure 5 B).

Since WRKYs are themselves regulated via phosphorylation by calcium-dependent protein kinases (CDPKs), and AM fungal colonisation upregulated calcium-binding proteins, we also examined calcium signalling components (Figure 5 C). Several CDPKs, calmodulin, and calmodulin-like genes were upregulated in AM fungi-colonised healthy plants.

To investigate how AM fungi influence defence-related hormone signalling, we analysed transcriptional changes in the ethylene, SA, and JA pathways. In healthy AM fungi-colonised plants, ethylene-responsive genes such as *EIN3* were upregulated, while defence-associated ethylene responses remained unchanged (Figure S3), indicating a shift towards metabolic rather than immune regulation. In contrast, several JA synthesis pathway genes, including *LOX* (lipoxygenase) genes, were expressed at lower levels in AM fungal-colonised diseased plants (AP) than in plants infected with *P. hordei* alone (P). This did not, however, affect downstream JA-responsive TIFY genes, which suggests that the effect on JA-synthesis genes can be unlinked from JA-signalling in this context (Figure S4).

Similarly, for the SA-biosynthesis pathway, AM fungi caused downregulation of CM (Chorismate mutase) and most PAL (Phenylalanine ammonia lyase) genes during *P. hordei* infection (Figure S4), while enhancing SA-mediated signalling, as seen by the induction of SA-responsive PR genes (Figure 3). Expression of *NPR1*, the central SA co-activator, was also upregulated in AM fungal plants but suppressed by *P. hordei* regardless of AM fungal status (Figure S4). In contrast, the strong induction of *PR* genes appears to be largely independent of *NPR1* transcription, which is not markedly affected by pathogen challenge. (Skelly, Frungillo and Spoel 2016; Withers and Dong 2016; Skelly *et al*. 2019). Overall, AM fungi reshaped transcriptional profiles of hormone pathways.

### AM fungi modulated expression of ubiquitin machinery genes without altering global ubiquitin levels

Gene Ontology enrichment analysis of AM fungal-induced transcripts showed enrichment of genes involved in protein modification processes (Figure 4 B), including ubiquitination-related genes. To determine whether this transcriptional signal translated into changes at the protein level, we assessed global protein ubiquitination across treatments (Figure S5). Western blot analysis showed that ubiquitin signal was significantly increased by *Puccinia hordei* infection (F*_1,8_* = 24.20, p = 0.001), with no significant effects of AM fungal inoculation or AM fungi × *Puccinia* interaction (Figure 6 A), indicating that AM fungi do not induce broad changes in total protein ubiquitination.

**Figure 6.**
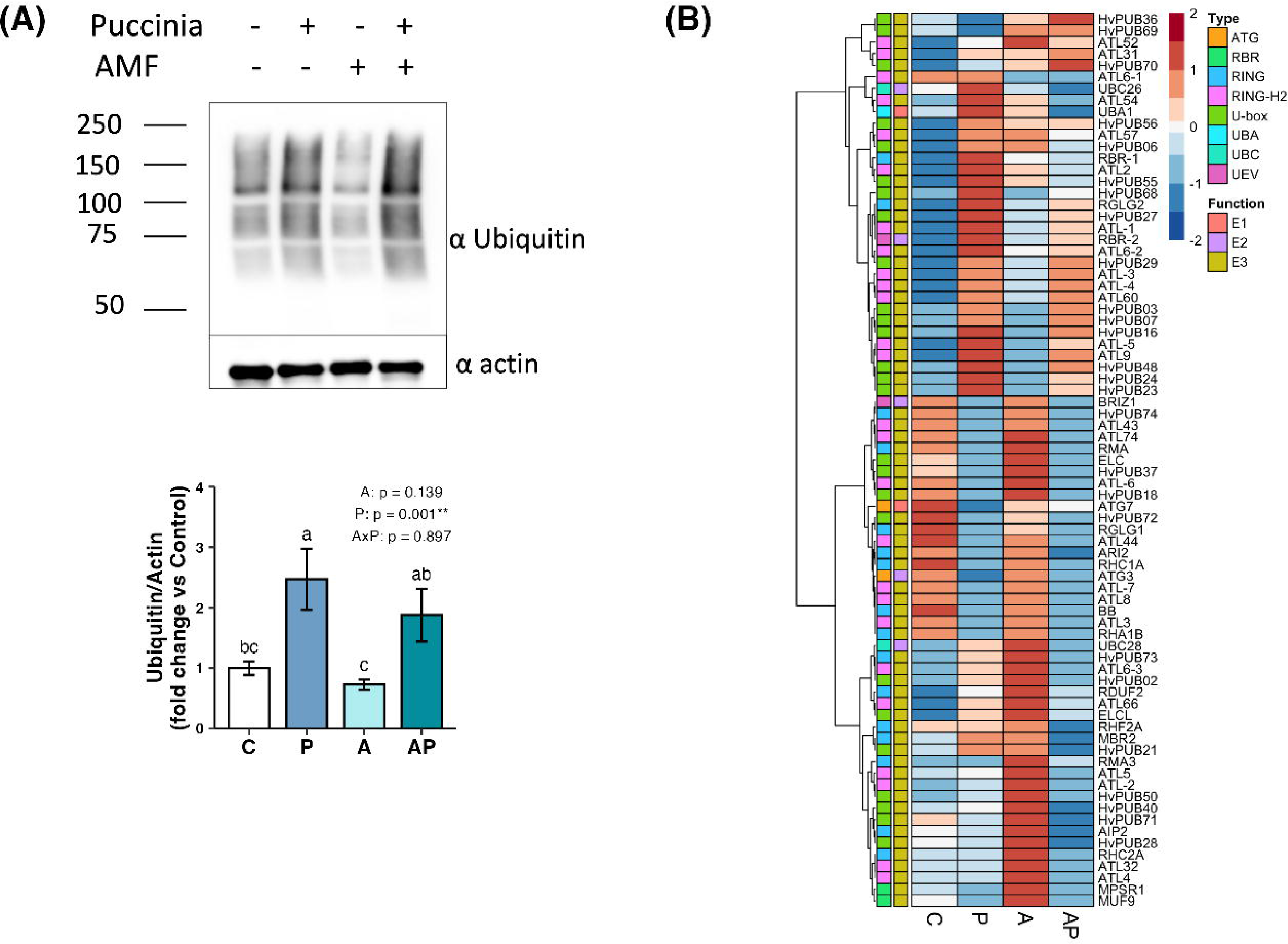
Effects of AMF and *Puccinia hordei* on ubiquitination. (A) Ubiquitinated protein levels normalised to actin and expressed relative to control (mean ± SE, n = 3); letters denote Tukey HSD groupings (P < 0.05) and ANOVA P-values derive from a two-way ANOVA on log₂-transformed fold-change values. (B) Row-scaled expression of differentially expressed ubiquitin-related genes across treatments, annotated by ubiquitination function and pathway type. Genes are annotated by function and ubiquitin pathway type, with PUB genes classified according to Ryu et al. (2019) and closest Arabidopsis homologues. Treatments are ordered as C (control), P (*P. hordei*), A (AMF), and AP (AMF + *P. hordei*) for consistency with the Western Blot.

Despite the absence of a global effect, transcriptomic analysis revealed substantial AM fungi-associated modulation of genes encoding ubiquitination enzymes. Many E3 ubiquitin ligase genes were differentially expressed, as well as some ubiquitin E1 (activating enzymes) and E2 (conjugating enzymes) (Figure 6 B). Expression patterns clustered into five distinct groups with the top cluster upregulated by AM fungi in healthy and infected plants (Figure 6 B). This small group included several RING/U-box-type E3 ligases from the ATL family and PUB families, suggesting that AM fungi selectively modulate specific components of the ubiquitin-proteasome system (UPS), rather than broadly altering the cellular ubiquitome. In fact, the largest cluster of ubiquitination-related differentially expressed genes (DEGs) was downregulated in AM-colonised plants during *P. hordei* infection (Figure 6 B), supporting a targeted rather than global modulation of ubiquitin signalling.

## Discussion

AM fungi have been shown to induce resistance against various pathogens of crop plants (Fiorilli *et al*. 2011, 2018; Campos-Soriano, García-Martínez and Segundo 2012), and therefore their application as an alternative crop protection method has been widely explored (Brito, Carvalho and Goss 2021; Lutz *et al*. 2023). In this study, we investigated whether colonisation by the arbuscular mycorrhizal (AM) fungus *Rhizophagus irregularis* influences barley fitness and disease outcomes during infection with the biotrophic foliar pathogen *Puccinia hordei*, within the broader context of sustainable crop protection and UN Sustainable Development Goal 2: Zero Hunger.

### AM fungal colonisation induced pathogen-independent transcriptional reprogramming

While AM colonisation did not result in significant growth promotion, reduced infection load, or mitigated *P. hordei*-induced biomass loss, it triggered extensive transcriptional changes in the host. Notably, many of these changes were observed in the absence of pathogen challenge, indicating that AM fungi substantially reshape host regulatory networks independently of disease pressure.

A range of studies have shown that inoculation with AM fungi can promote plant growth, grain yield and vigour (Pellegrino *et al*. 2015; Fiorilli *et al*. 2018; Lutz *et al*. 2023). However, in our study, *Rhizophagus irregularis* colonisation did not result in measurable growth promotion under the tested conditions. This absence of a growth response may reflect limited compatibility between the specific host-symbiont combination, as plant responses to AM fungi are known to vary widely depending on both the fungal isolate and the host genotype (Munkvold *et al*. 2004; Hong *et al*. 2012; Koch *et al*. 2017). Barley cultivars exhibit varying degrees of mycorrhizal responsiveness, with differences in colonisation efficiency and functional outcomes (Peña Venegas *et al*. 2021; Gebremeskel *et al*. 2024; Marrassini *et al*. 2024). These genotype-dependent effects likely contribute to the observed lack of growth benefit in our system and suggest that the *Hordeum vulgare* cv. Bowman – *R. irregularis* pairing forms a transcriptionally active symbiosis with limited physiological output under the tested conditions.

In pathogen-free plants, AM colonisation has been shown to induce extensive transcriptional reprogramming associated with symbiosis establishment and maintenance. These changes include broad remodelling of primary metabolism and carbon allocation, reflecting a shift in host metabolic priorities rather than activation of defence responses (Smith and Smith 2011). Such transcriptional adjustments are thought to support symbiotic function and signalling while avoiding the fitness costs associated with constitutive immunity. Importantly, numerous studies have shown that these pathogen-independent molecular changes can occur without measurable penalties to host growth, highlighting that AM fungi can markedly reshape host regulatory and metabolic states while maintaining overall plant fitness (Smith and Smith 2011; Koch *et al*. 2017).

### Pathogen-dependent immune priming in AM-colonised barley

In contrast to the pathogen-independent effects described above, a distinct set of transcriptional responses emerged specifically following *P. hordei* infection. Despite the absence of reduced pathogen burden, AM-colonised plants displayed enhanced induction of genes involved in broad-spectrum immune responses, including salicylic acid (SA)-associated pathways and cell wall defence mechanisms. Such patterns are consistent with immune priming, a hallmark of mycorrhiza-induced resistance (MIR), in which defence responses are sensitised but not constitutively activated (Jung *et al*. 2012; Huot *et al*. 2014).

Crucially, priming does not always confer full immunity. Instead, it can result in partial resistance, slower pathogen development, or reduced sporulation, outcomes that are often not captured by static disease assessments (Pieterse *et al*. 2014). This aligns with previous studies showing that MIR often manifests as subtle, ecologically relevant phenotypes such as delayed symptom onset or attenuated disease progression (Pozo *et al*. 2002; Pozo and Azcón-Aguilar 2007; Jung *et al*. 2012). Future experiments quantifying disease dynamics over time (e.g. latent period, spore production) could reveal benefits of MIR that static measurements miss.

Whether broad-spectrum defence priming alone is sufficient for MIR establishment remains debated. Several studies have demonstrated that AM fungi can induce pathogen-specific immune reprogramming distinct from that of non-colonised plants (Campos-Soriano, García-Martínez and Segundo 2012; Fiorilli *et al*. 2018). In our study, AM fungal colonisation attenuated the overall magnitude of the *P. hordei*-induced transcriptional response and led to the activation of a specific subset of defence-related genes. However, this reprogramming alone was insufficient to confer effective resistance against a highly specialised biotrophic pathogen such as *P. hordei*. As MIR is highly context-dependent and often biases JA- and ethylene- mediated defences, it can antagonise SA pathway responses important against biotrophic pathogens (Cameron *et al*. 2013; Liu *et al*. 2023). This could be reflected in the elevated defence-related transcript levels, as seen here, without a corresponding increase in downstream immune outputs (Caldo, Nettleton and Wise 2004; Zierold, Scholz and Schweizer 2005; Polesani *et al*. 2010). Moreover, pathogen effectors can bypass the priming by targeting specific immune proteins or disturbing host proteostasis (Bauters, Stojilković and Gheysen 2021; Kainat *et al*. 2025). Therefore, our findings suggest that the outcome of AM colonisation in the context of pathogen attack is not solely determined by transcriptional priming but likely emerges from a complex tripartite interaction between host plant, mycorrhizal symbiont, and pathogen. Disentangling these multilayered interactions remains a significant challenge and will require more refined tools and approaches to fully understand the context-dependence of mycorrhiza-induced resistance.

### Transcriptional and post-translational regulators of the primed immune state

Our data identify WRKY transcription factors as potential regulators of the primed immune state induced by *R. irregularis* in barley. Remarkably, nearly half of all annotated WRKY genes were differentially expressed, representing a broader-scale alteration of regulatory networks than previously reported in other systems such as potato, tomato, or apple (Gallou, Declerck and Cranenbrouck 2012; Cervantes-Gámez *et al*. 2016; Wang *et al*. 2022). This widespread regulation suggests that AM fungi fine-tune immune responses and developmental trade-offs through WRKY-mediated transcriptional control. The co-induction of calcium-dependent protein kinases and calmodulin-related genes further supports the involvement of calcium-WRKY signalling modules in MIR, consistent with findings in other host-AM fungal systems (Campos-Soriano, García-Martínez and Segundo 2012; Fiorilli *et al*. 2018).

In parallel, AM colonisation altered the expression of hormonal signalling genes in a manner that deviates from canonical models of MIR. Suppression of jasmonic acid (JA) biosynthesis genes and upregulation of TIFY repressors, which inhibit JA signalling, indicates a shift away from typical JA/ethylene-mediated responses. Interestingly, while salicylic acid (SA) biosynthesis and signalling genes were downregulated, SA-responsive defence markers were upregulated during pathogen challenge. This pattern is consistent with an immune priming mechanism previously described in rice, where AM fungi induced expression of *NPR1*, the central coactivator of SA-responsive gene expression, in the leaves of healthy colonised plants without activating the PR gene expression (Campos-Soriano and Segundo 2011).

This apparent disconnect between NPR1 transcript levels and downstream defence activation is consistent with previous findings that *NPR1* activity is predominantly regulated at the post-translational level through redox-dependent conformational changes, phosphorylation and proteasomal turnover, rather than transcriptional control alone (Skelly, Frungillo and Spoel 2016; Withers and Dong 2016; Skelly *et al*. 2019). These findings suggest that AM fungi establish a preconditioned immune state that can be rapidly mobilised upon pathogen challenge yet may be insufficient to overcome the virulence strategies of highly specialised biotrophic pathogens such as *P. hordei*. Such priming is therefore likely to depend not only on transcriptional readiness, but also on post-translational regulatory mechanisms governing immune protein activation and stability.

In addition to transcriptional reprogramming, we observed significant modulation of the host ubiquitin-proteasome system (UPS). Although global levels of ubiquitinated proteins remained unchanged, AM colonisation led to the upregulation of numerous components of the ubiquitination machinery, including E3 ligases from the PUB and ATL families (Figure 6 B), which are known to regulate diverse stress responses (Guzmán 2012; Trujillo 2018). This seemingly paradoxical pattern may reflect increased protein turnover and dynamic post-translational regulation of immune signalling, rather than broad immune activation. Notably, PUB and ATL ligases have established roles in regulating stress and immune responses, including chitin-triggered signalling and callose deposition (Berrocal-Lobo *et al*. 2010; Maekawa *et al*. 2012). Together, our findings suggest that AM fungi-induced immune priming operates through a multilayered regulatory framework, integrating transcriptional, hormonal, and post-translational controls, to enhance defence readiness while maintaining metabolic balance.

## Conclusion

Achieving global food security under increasing biotic stress requires sustainable strategies that reduce crop losses while minimising chemical inputs. While AM fungal colonisation did not reduce pathogen burden at the phenotypic level in this study, the extensive transcriptional reprogramming points to a robust immune priming effect that may confer delayed or context-dependent protection not captured within the experimental timeframe of this study. Importantly, this immune activation occurred without measurable growth penalties, suggesting that *R. irregularis,* or other AM fungi, could, pending field validation, serve as a sustainable tool to enhance crop resilience.

Realising this potential will require integrating AM fungal strategies into broader crop protection frameworks and continued research into the ecological and mechanistic dependencies that shape mycorrhiza-induced resistance. By leveraging the beneficial properties of symbiotic soil fungi, we may unlock new ways to support plant health and productivity in alignment with the goals of Zero Hunger and sustainable agriculture.

## Data availability

RNA-sequence data generated in this study have been deposited in the NCBI Sequence Read Archive (SRA) under accession number PRJNA1328502. The data and R code that support the findings of this study are openly available as a GitHub repository, archived at https://zenodo.org/records/18312009.

