## Supplementary data for "AMF primes immune genes against *Puccinia hordei* (Brown rust) in *Hordeum vulgare* but does not reduce pathogen burden"

Supplementary data containing four supplementary tables (S1–S4) with further information about qPCR primers (Table S1), RNA sequencing summary statistics (Table S2), differentially expressed genes in AMF inoculated plants (Table S3) and differentially expressed WRKY transcription factors (Table S4).

Additional figures provide information on barley biomass allocation (Figure S1), the number of up- and down-regulated genes in each treatment combination (Figure S2), Ethylene signalling gene expression (Figure S3), J**asmonic acid (JA) and salicylic acid (SA) signalling gene expression (Figure S4), and the Ubiquitin Western blot (Figure S5).**

Table S1 Primers used in qPCR assays.

| **Target** | **Sequence (**5′–3′**)** | **Reference** |
| --- | --- | --- |
| *HvACT* | GCCGTGCTTTCCCTCTATG  GAAGGAGTAACCTCTCTCGG | This study |
| *HvPR1b* | GCTAGCCATCTTGCTCGCC  GCTTGCAGTCGTTGATCCTC | Adapted from (Gao *et al.* 2018) |
| *HvPR2* | GATGTTGCCTCCATGTTTGCAG  GCATGCCGTTGATGCCCTTG | Adapted from (Gao *et al.* 2018) |
| *HvPR3* | GTTCCAGGCTACGGTGTAATC  GTTCCGTTGGGTGTAGCAGT | Adapted from (Gao *et al.* 2018) |
| *WRKY28* | CATGTGTTTCAACCCGTTCCAG  GAAGGCAGAAATGTCGAAGTTGG | Adapted from (Meng and Wise 2012) |
| *PhEF2* | CCCTGAAAATGCTTTGGGTGG  TTGACGGTGTACATGGGAGTG | This study |
| AMF18S | ATAGGGATAGTTGGGGGCAT  GTTTCCCGTAAGGCGCCGAA | (Helgason *et al.* 1998; Hewins, Carrino-Kyker and Burke 2015) |
| *HvEF1*α | GAAGATGATTCCCACCAAGC  TGACACCAACAGCCACAGTT | (Hua *et al.* 2015) |

Table S2 Summary statistics of RNA sequencing data from three libraries from C, control; A, AMF addition; P, Puccinia addition; AP, AMF plus Puccinia addition treatments.

| **Sample name** | **Raw reads (millions)** | **Clean reads (millions)** | **Total clean bases (Gb)** | **GC (%)** |
| --- | --- | --- | --- | --- |
| C-1 | 40.74 | 39.69 | 5.95 | 54 |
| C-2 | 53.65 | 52.14 | 7.82 | 55 |
| C-3 | 40.44 | 39.24 | 5.89 | 55 |
| A-1 | 44.47 | 42.89 | 6.43 | 55 |
| A-2 | 42.96 | 41.52 | 6.23 | 56 |
| A-3 | 41.07 | 39.77 | 5.96 | 55 |
| P-1 | 8.18 | 7.84 | 1.18 | 52 |
| P-2 | 44.42 | 42.78 | 6.42 | 50 |
| P-3 | 46.12 | 44.40 | 6.66 | 51 |
| AP-1 | 43.78 | 42.42 | 6.36 | 51 |
| AP-2 | 44.04 | 42.65 | 6.40 | 49 |
| AP-3 | 44.52 | 43.12 | 6.47 | 51 |

C, control, plants not inoculated or infected; A, plants inoculated with the AM fungus R. irregularis; P, plants infected with the pathogen P. hordei; A+P, plants first inoculated with the AM fungus R. irregularis and then infected with the pathogen P. hordei 28 days later. RNA analysis was conducted on barley leaves 7 dpi.

Table S3 Differentially expressed genes in control Vs AMF inoculated plants.

| **Gene ID (HORVU.MOREX.r3.)** | **Protein ID** | **Description** | **logFC** | **logCPM** | **FDR** |
| --- | --- | --- | --- | --- | --- |
| 5HG0497680 | Q3T5P5 | C-repeat binding factor 6 (CBF6) (CRT/DRE binding factor 6) | 3.09 | 5.00 | 1.16E-22 |
| 3HG0321580 | A0A8I6X9M1 | Uncharacterized protein | 3.05 | 4.43 | 9.85E-19 |
| 3HG0323580 | A0A8I7BB67 | Uncharacterized protein | 3.03 | 5.10 | 2.55E-22 |
| 1HG0020440 | A0A8I7B2J6 | Protein kinase domain-containing protein | 2.98 | 3.33 | 4.39E-11 |
| 2HG0187050 | A0A8I6WYA0 | Protein DETOXIFICATION (Multidrug and toxic compound extrusion protein) | 2.95 | 3.73 | 3.15E-13 |
| 6HG0603700 | Q8S4R3 | CRT/DRE binding factor 1 (HvCBF1) | 2.92 | 4.30 | 7.13E-17 |
| 7HG0743270 | A0A8I6YNK5 | WRKY domain-containing protein | 2.86 | 3.34 | 2.10E-10 |
| 7HG0743280 | A0A8I6YK40 | WRKY domain-containing protein | 2.73 | 7.09 | 7.32E-18 |
| 7HG0719250 | A0A8I6Z6J5 | RING-type E3 ubiquitin transferase (EC 2.3.2.27) | 2.71 | 4.10 | 1.72E-13 |
| 2HG0109800 | A0A8I6WGM7 | J domain-containing protein | 2.59 | 2.61 | 2.78E-06 |
| 7HG0645900 | F2EDZ5 | Predicted protein | 2.58 | 3.70 | 1.73E-10 |
| 4HG0406580 | F2E7C4 | Predicted protein | 2.58 | 3.25 | 1.95E-08 |
| 1HG0057750 | M0YM29 | Uncharacterized protein | 2.55 | 3.95 | 1.63E-11 |
| 7HG0645780 | A0A8I6YAU9 | Uncharacterized protein | 2.55 | 3.53 | 1.66E-09 |
| 7HG0668940 | F2D820 | U-box domain-containing protein (EC 2.3.2.27) (RING-type E3 ubiquitin transferase PUB) | 2.52 | 6.32 | 5.94E-17 |
| 2HG0192970 | A0A8I6WX32 | VQ domain-containing protein | 2.52 | 2.65 | 3.94E-06 |
| 6HG0619310 | A0A8I7BHJ9 | AP2/ERF domain-containing protein | 2.51 | 4.37 | 3.12E-13 |
| 3HG0322130 | A0A8I6Y2K4 | EF-hand domain-containing protein | 2.49 | 2.88 | 1.19E-06 |
| 6HG0598010 | A0A8I7BH45 | AP2/ERF domain-containing protein | 2.49 | 3.35 | 2.48E-08 |
| 2HG0186680 | A0A8I6WV37 | H15 domain-containing protein | 2.46 | 2.01 | 1.51E-04 |
| 6HG0600160 | F2DWS3 | Predicted protein | 2.45 | 2.19 | 7.75E-05 |
| 7HG0643210 | F2EHR1 | Predicted protein | 2.43 | 4.38 | 1.84E-12 |
| 7HG0669300 | A0A8I6Z8V8 | Protein TIFY (Jasmonate ZIM domain-containing protein) | 2.42 | 2.97 | 1.23E-06 |
| 4HG0406650 | F2EL34 | Predicted protein | 2.38 | 4.06 | 9.21E-11 |
| 4HG0338620 | A0A8I6WYF0 | Uncharacterized protein | 2.34 | 2.69 | 1.56E-05 |
| 2HG0176250 | F2DJB1 | Predicted protein | 2.33 | 2.44 | 5.69E-05 |
| 2HG0208110 | A0A8I7B8E7 | Uncharacterized protein | 2.33 | 3.36 | 1.82E-07 |
| 3HG0222590 | A0A8I6X3A8 | Patatin (EC 3.1.1.-) | 2.27 | 3.33 | 4.78E-07 |
| 2HG0159040 | A0A8I6WNN4 | Transcription factor (bHLH transcription factor) (Basic helix-loop-helix protein) | 2.25 | 3.09 | 3.55E-06 |
| 6HG0546340 | F2DNU8 | Peroxidase (EC 1.11.1.7) | 2.24 | 5.60 | 8.14E-14 |
| 4HG0340040 | F2DFR9 | Predicted protein | 2.24 | 3.27 | 1.17E-06 |
| 4HG0383400 | F2CZR7 | Predicted protein (Putative roothairless 3) | 2.21 | 6.23 | 2.34E-13 |
| 1HG0080800 | A0A8I7B3Y3 | [RNA-polymerase]-subunit kinase (EC 2.7.11.23) | 2.16 | 4.05 | 5.42E-09 |
| 4HG0406630 | F2E0Z0 | Predicted protein | 2.14 | 4.07 | 6.12E-09 |
| 3HG0239530 | A0A8I7B9B0 | Uncharacterized protein | 2.13 | 6.99 | 1.59E-11 |
| 3HG0276810 | A0A8I6X8J6 | WRKY domain-containing protein | 2.11 | 5.65 | 2.40E-12 |
| 3HG0246250 | A0A8I6XWS0 | Fe2OG dioxygenase domain-containing protein | 2.11 | 2.22 | 8.17E-04 |
| 6HG0605940 | A0A8I6Y6A9 | Xyloglucan endotransglucosylase/hydrolase (EC 2.4.1.207) | 2.08 | 5.81 | 4.32E-12 |
| 3HG0321590 | A0A8I6Y739 | BHLH domain-containing protein | 2.05 | 2.75 | 1.50E-04 |
| 7HG0744280 | A0A8I6ZFB6 | Uncharacterized protein | 2.04 | 3.79 | 2.90E-07 |
| 6HG0568570 | F2D8Q7 | Predicted protein | 2.04 | 7.49 | 6.56E-10 |
| 3HG0313560 | F2EEZ3 | Predicted protein | 2.03 | 2.23 | 1.28E-03 |
| 1HG0057580 | A0A8I6WTR7 | PRA1 family protein | 2.03 | 3.07 | 3.68E-05 |
| 3HG0291910 | F2CTB8 | Peroxidase (EC 1.11.1.7) | 2.02 | 2.59 | 3.82E-04 |
| 2HG0212840 | A0A8I6X0L7 | GST N-terminal domain-containing protein | 1.99 | 3.29 | 1.63E-05 |
| 5HG0513730 | A0A8I7BFF6 | Formin-like protein | 1.99 | 5.64 | 3.81E-11 |
| 2HG0111820 | F2CUU0 | Predicted protein | 1.99 | 5.07 | 1.72E-10 |
| 6HG0617580 | A0A8I6Y7L2 | AP2/ERF domain-containing protein | 1.99 | 2.42 | 9.21E-04 |
| 5HG0430240 | A0A287QCK8 | E3 ubiquitin-protein ligase RMA (EC 2.3.2.27) (Protein RING membrane-anchor) (RING-type E3 ubiquitin transferase RMA) | 1.98 | 4.53 | 3.43E-09 |
| 6HG0560670 | F2D8F9 | Predicted protein | 1.97 | 4.05 | 1.27E-07 |
| 1HG0005420 | A0A8I6W3V3 | Glycosyltransferase 61 catalytic domain-containing protein | 1.97 | 6.82 | 3.55E-10 |
| 3HG0297880 | A0A8I7BAK1 | Ubiquitin-like domain-containing protein | 1.96 | 2.48 | 9.42E-04 |
| 3HG0302610 | M0Z904 | Uncharacterized protein | 1.94 | 2.07 | 3.85E-03 |
| 2HG0213970 | F2DRK3 | poly(A)-specific ribonuclease (EC 3.1.13.4) | 1.94 | 6.29 | 1.91E-10 |
| 7HG0732170 | A0A8I6ZEA3 | EF-hand domain-containing protein | 1.94 | 6.37 | 2.21E-10 |
| 5HG0474120 | M0W8U1 | Uncharacterized protein | 1.93 | 7.16 | 2.53E-09 |
| 5HG0509280 | M0X426 | Uncharacterized protein | 1.92 | 4.20 | 9.96E-08 |
| 2HG0139040 | A0A8I6WJ92 | Uncharacterized protein | 1.92 | 4.49 | 1.57E-08 |
| 3HG0322070 | A0A8I6XCA7 | EF-hand domain-containing protein | 1.90 | 3.61 | 7.29E-06 |
| 5HG0437580 | A0A8I6Y5R5 | Uncharacterized protein | 1.90 | 3.35 | 3.08E-05 |
| 7HG0660720 | A0A8I6Z0F2 | RING-type E3 ubiquitin transferase (EC 2.3.2.27) | 1.89 | 5.85 | 3.37E-10 |
| 1HG0090520 | F2E0Y3 | Predicted protein | 1.89 | 4.00 | 7.38E-07 |
| 7HG0653970 | A0A8I6Z792 | Uncharacterized protein | 1.87 | 3.39 | 3.66E-05 |
| 5HG0484180 | F2DT38 | Predicted protein | 1.85 | 6.25 | 1.21E-09 |
| 2HG0181540 | M0YYK1 | AAA+ ATPase domain-containing protein | 1.84 | 5.14 | 3.36E-09 |
| 2HG0198540 | A0A8I6XIB5 | Uncharacterized protein | 1.83 | 2.54 | 2.18E-03 |
| 1HG0072150 | A0A8I6WSB6 | Uncharacterized protein | 1.81 | 3.04 | 3.48E-04 |
| 1HG0079460 | A0A8I7B1H5 | Protein kinase domain-containing protein | 1.81 | 3.75 | 9.42E-06 |
| 6HG0574290 | A0A8I6YTS4 | Uncharacterized protein | 1.80 | 5.36 | 4.60E-09 |
| 3HG0283700 | A0A8I6XPX1 | Transcription factor (bHLH transcription factor) (Basic helix-loop-helix protein) | 1.79 | 3.75 | 1.30E-05 |
| 2HG0127480 | A0A8I6XEA2 | NAD-dependent epimerase/dehydratase domain-containing protein | 1.78 | 2.32 | 5.35E-03 |
| 4HG0383630 | A0A8I7B7A5 | Wound-induced protein 1 | 1.77 | 3.88 | 7.55E-06 |
| 1HG0069640 | F2DTF4 | Predicted protein | 1.77 | 2.27 | 6.73E-03 |
| 7HG0744270 | A0A8I6YK73 | Uncharacterized protein | 1.77 | 3.84 | 1.00E-05 |
| 7HG0709950 | A0A8I6YQB9 | HMA domain-containing protein | 1.77 | 2.14 | 9.44E-03 |
| 3HG0274050 | A0A8I6XNM2 | Uncharacterized protein | 1.76 | 3.11 | 4.67E-04 |
| 1HG0090460 | A0A8I6X7C3 | WRKY domain-containing protein | 1.73 | 5.05 | 4.63E-08 |
| 2HG0122190 | A0A8I6WYG2 | RING-type E3 ubiquitin transferase (EC 2.3.2.27) | 1.73 | 2.17 | 1.11E-02 |
| 1HG0080940 | F2CRY4 | Predicted protein (WRKY transcription factor 19) | 1.73 | 3.25 | 3.37E-04 |
| 5HG0512730 | A0A8I6YVF6 | C2H2-type domain-containing protein | 1.73 | 7.10 | 1.27E-07 |
| 1HG0053590 | A0A8I6W639 | Lysine-specific demethylase JMJ706 | 1.72 | 5.45 | 2.08E-08 |
| 1HG0029360 | F2CZY9 | Predicted protein | 1.70 | 5.23 | 4.70E-08 |
| 5HG0429040 | A0A8I6XHU3 | VQ domain-containing protein | 1.70 | 2.23 | 1.11E-02 |
| 6HG0575580 | A0A8I6YAD0 | Transcription factor MYB44 | 1.69 | 2.32 | 9.48E-03 |
| 5HG0504260 | A0A287S5S4 | Auxin efflux carrier component | 1.69 | 3.67 | 6.93E-05 |
| 7HG0664030 | A0A8I6YKQ3 | Uncharacterized protein | 1.68 | 5.31 | 6.84E-08 |
| 3HG0287070 | A0A8I6XX25 | Glycosyltransferase (EC 2.4.1.-) | 1.67 | 4.38 | 2.29E-06 |
| 7HG0664780 | A0A8I6Z0T8 | Patatin (EC 3.1.1.-) | 1.67 | 2.92 | 2.12E-03 |
| 7HG0667900 | F2D2W1 | Predicted protein | 1.67 | 6.46 | 9.99E-08 |
| 5HG0526110 | F2EKK3 | Predicted protein | 1.67 | 2.79 | 3.20E-03 |
| 2HG0123480 | A0A8I6WPV6 | WRKY domain-containing protein | 1.66 | 7.93 | 2.83E-06 |
| 1HG0071500 | A0A8I6WHP1 | WRKY domain-containing protein | 1.65 | 5.16 | 1.78E-07 |
| 4HG0409510 | A0A8I6YKG1 | NAC domain-containing protein | 1.63 | 4.55 | 2.27E-06 |
| 5HG0527250 | A0A8I6YE52 | Bowman-Birk serine protease inhibitors family domain-containing protein | 1.63 | 3.24 | 9.06E-04 |
| 5HG0480150 | F2E2B3 | Predicted protein | 1.63 | 2.37 | 1.28E-02 |
| 7HG0751680 | A0A8I6YPM6 | NAC domain-containing protein | 1.63 | 6.25 | 1.58E-07 |
| 5HG0460570 | M0V1H8 | Glycosyltransferase (EC 2.4.1.-) | 1.62 | 5.89 | 1.27E-07 |
| 2HG0113890 | M0XJ21 | Indole-3-acetic acid-amido synthetase GH3.9 | 1.62 | 2.55 | 8.90E-03 |
| 2HG0204690 | A0A8I7B8A4 | Uncharacterized protein | 1.62 | 3.24 | 9.43E-04 |
| 2HG0122140 | A0A8I6XDQ2 | EF-hand domain-containing protein | 1.62 | 6.40 | 2.28E-07 |
| 1HG0021970 | A0A8I6WNA0 | Fe2OG dioxygenase domain-containing protein | 1.62 | 9.36 | 6.24E-05 |
| 2HG0212050 | A0A8I6XID2 | Receptor-like serine/threonine-protein kinase (EC 2.7.11.1) | 1.61 | 3.52 | 3.18E-04 |
| 3HG0240670 | F2CUP3 | Histone H4 | 1.61 | 2.08 | 2.45E-02 |
| 3HG0331020 | A0A8I6XDW8 | C3H1-type domain-containing protein | 1.61 | 9.54 | 9.11E-05 |
| 2HG0099170 | A0A8I6WZY7 | NADH:flavin oxidoreductase/NADH oxidase N-terminal domain-containing protein | 1.60 | 2.35 | 1.52E-02 |
| 7HG0668680 | A0A8I6YDJ0 | Hexosyltransferase (EC 2.4.1.-) | 1.60 | 5.78 | 1.84E-07 |
| 1HG0055700 | A0A8I6WTZ3 | RING-type E3 ubiquitin transferase (EC 2.3.2.27) | 1.60 | 2.12 | 2.44E-02 |
| 3HG0237810 | F2CX90 | Predicted protein | 1.60 | 3.59 | 2.94E-04 |
| 4HG0406620 | A0A8I6XTW7 | Uncharacterized protein | 1.59 | 3.44 | 5.64E-04 |
| 4HG0392530 | A0A8I7BCP8 | Thaumatin-like protein 1 | 1.59 | 2.57 | 1.03E-02 |
| 6HG0622710 | A0A8I6Y881 | Dehydrin 5 | 1.58 | 2.04 | 3.05E-02 |
| 6HG0574180 | A0A8I6Y3R5 | Uncharacterized protein | 1.58 | 2.58 | 1.04E-02 |
| 5HG0462600 | A0A8I6XB33 | Receptor-like serine/threonine-protein kinase (EC 2.7.11.1) | 1.58 | 3.82 | 1.23E-04 |
| 3HG0278170 | F2DI80 | Predicted protein | 1.58 | 4.07 | 3.91E-05 |
| 7HG0650970 | F2CQJ8 | Histone H3 | 1.58 | 2.59 | 1.05E-02 |
| 7HG0711070 | A0A8I6ZBQ0 | Protein kinase domain-containing protein | 1.57 | 4.49 | 6.88E-06 |
| 2HG0200920 | A0A8I6XDE5 | Agmatine coumaroyltransferase-2 | 1.57 | 2.93 | 4.32E-03 |
| 3HG0310480 | A0A8I6XAT4 | Nucleotide-diphospho-sugar transferase domain-containing protein | 1.56 | 4.71 | 3.67E-06 |
| 1HG0089070 | F2CZB9 | Predicted protein | 1.55 | 3.27 | 1.64E-03 |
| 4HG0414340 | A0A8I6Y3C7 | Uncharacterized protein | 1.55 | 6.59 | 1.28E-06 |
| 1HG0010340 | F2DLF7 | Predicted protein | 1.55 | 2.39 | 1.90E-02 |
| 2HG0212970 | A0A8I6WV53 | Phospholipase/carboxylesterase/thioesterase domain-containing protein | 1.54 | 3.12 | 2.93E-03 |
| 3HG0320050 | M0XJI3 | EF-hand domain-containing protein | 1.54 | 6.46 | 1.17E-06 |
| 7HG0669660 | F2D229 | Predicted protein | 1.54 | 4.66 | 5.97E-06 |
| 3HG0225490 | A0A8I7B4M1 | Uncharacterized protein | 1.54 | 2.72 | 9.95E-03 |
| 4HG0403480 | F2CZR1 | Predicted protein | 1.54 | 6.07 | 8.34E-07 |
| 7HG0732710 | A0A8I6YIP0 | DUF642 domain-containing protein | 1.53 | 2.80 | 8.68E-03 |
| 4HG0333220 | A0A8I6XL70 | WRKY domain-containing protein | 1.53 | 4.90 | 3.25E-06 |
| 5HG0426410 | F2DMI5 | Predicted protein | 1.52 | 5.19 | 1.80E-06 |
| 7HG0686980 | A0A8I6YEU8 | HSF-type DNA-binding domain-containing protein | 1.52 | 3.75 | 3.30E-04 |
| 2HG0160240 | F2CQC9 | Exocyst subunit Exo70 family protein | 1.52 | 5.74 | 1.04E-06 |
| 2HG0193650 | F2DND7 | Ubiquinol oxidase (EC 1.10.3.11) | 1.51 | 5.69 | 1.17E-06 |
| 4HG0401170 | Q70DK2 | Blue copper binding protein | 1.51 | 6.58 | 2.41E-06 |
| 1HG0016710 | F2CQV6 | Annexin | 1.51 | 3.90 | 2.00E-04 |
| 2HG0195500 | A0A8I6XHH4 | Uncharacterized protein | 1.50 | 2.40 | 2.45E-02 |
| 2HG0108680 | A0A8I7B249 | Methyltransferase type 11 domain-containing protein | 1.50 | 5.33 | 2.06E-06 |
| 3HG0302230 | F2DS14 | Predicted protein | 1.50 | 5.48 | 1.73E-06 |
| 5HG0524540 | F6MEL2 | Stress-induced transcription factor SNAC1 | 1.50 | 8.65 | 9.93E-05 |
| 2HG0194350 | A0A8I6WX71 | Uncharacterized protein | 1.50 | 2.91 | 8.14E-03 |
| 5HG0487040 | A0A8I6YSU8 | Alpha/beta hydrolase fold-3 domain-containing protein | 1.50 | 2.01 | 4.82E-02 |
| 5HG0478850 | F2DQH1 | Trehalose 6-phosphate phosphatase | 1.49 | 2.21 | 3.60E-02 |
| 3HG0229400 | A0A8I7B922 | Protein kinase domain-containing protein | 1.49 | 3.19 | 3.73E-03 |
| 3HG0319350 | A0A8I7BB31 | Annexin | 1.49 | 5.25 | 2.78E-06 |
| 4HG0386490 | A0A8I6XIX3 | Protein kinase domain-containing protein | 1.49 | 5.74 | 1.84E-06 |
| 2HG0119830 | F2CSE1 | 4-coumarate--CoA ligase (EC 6.2.1.12) | 1.48 | 2.86 | 1.03E-02 |
| 1HG0091390 | A0A8I6W8H9 | EF-hand domain-containing protein | 1.48 | 4.75 | 1.11E-05 |
| 2HG0126460 | A0A8I6WSH7 | Sulfotransferase (EC 2.8.2.-) | 1.48 | 2.19 | 3.97E-02 |
| 6HG0574190 | A0A8I6YIM7 | Uncharacterized protein | 1.48 | 3.78 | 4.95E-04 |
| 2HG0196540 | A0A8I6X7A0 | Malectin-like domain-containing protein | 1.47 | 6.74 | 6.54E-06 |
| 3HG0301110 | A0A8I6X2I3 | Nematode resistance protein-like HSPRO2 | 1.47 | 8.10 | 6.78E-05 |
| 4HG0383820 | A0A8I7B7A6 | Uncharacterized protein | 1.47 | 2.58 | 2.10E-02 |
| 5HG0517140 | F2CYV6 | Predicted protein | 1.46 | 5.38 | 3.89E-06 |
| 7HG0744260 | A0A8I6YNP6 | Uncharacterized protein | 1.45 | 5.47 | 3.94E-06 |
| 5HG0523850 | A0A287SQ79 | UEV domain-containing protein | 1.45 | 4.04 | 2.21E-04 |
| 3HG0267770 | A0A8I6XMY7 | DDE Tnp4 domain-containing protein | 1.45 | 7.83 | 5.87E-05 |
| 4HG0413770 | A0A8I6XIQ6 | F-box domain-containing protein | 1.45 | 5.30 | 5.64E-06 |
| 2HG0213820 | F2DFZ2 | Predicted protein | 1.44 | 7.76 | 5.69E-05 |
| 5HG0518130 | F2DCN6 | Predicted protein | 1.44 | 3.98 | 3.37E-04 |
| 5HG0490490 | F2E6Z8 | Predicted protein | 1.44 | 6.90 | 1.46E-05 |
| 4HG0337740 | A0A8I6WYC6 | Protein kinase domain-containing protein | 1.44 | 4.06 | 2.58E-04 |
| 3HG0290880 | A0A8I6XPB5 | UDP-arabinopyranose mutase (EC 5.4.99.30) | 1.43 | 3.89 | 5.38E-04 |
| 4HG0380790 | A0A8I6Y7B3 | VQ domain-containing protein | 1.43 | 5.49 | 5.86E-06 |
| 3HG0300120 | A0A8I6XH61 | Late embryogenesis abundant protein LEA-2 subgroup domain-containing protein | 1.43 | 7.73 | 6.34E-05 |
| 5HG0469490 | F2DD32 | Predicted protein | 1.42 | 2.45 | 3.44E-02 |
| 7HG0724480 | A0A8I6ZDP3 | VQ domain-containing protein | 1.42 | 2.96 | 1.21E-02 |
| 6HG0590970 | F2D0B7 | Predicted protein | 1.42 | 6.51 | 1.14E-05 |
| 5HG0487980 | A0A8I6Y2S7 | RING-type E3 ubiquitin transferase (EC 2.3.2.27) | 1.41 | 4.36 | 1.12E-04 |
| 5HG0507470 | A0A8I6XT32 | Arogenate dehydratase (EC 4.2.1.91) | 1.41 | 4.81 | 2.92E-05 |
| 5HG0478330 | A0A8I6YRW2 | AN1-type domain-containing protein | 1.41 | 3.88 | 7.38E-04 |
| 6HG0618950 | F2DDS9 | Predicted protein | 1.40 | 6.42 | 1.30E-05 |
| 3HG0245610 | A0A8I7B517 | RING-type E3 ubiquitin transferase (EC 2.3.2.27) | 1.40 | 5.89 | 9.17E-06 |
| 7HG0657470 | A0A8I6YBX3 | Fe2OG dioxygenase domain-containing protein | 1.40 | 3.77 | 1.20E-03 |
| 7HG0640720 | A0A8I7BI27 | non-specific serine/threonine protein kinase (EC 2.7.11.1) | 1.40 | 5.83 | 9.17E-06 |
| 7HG0746660 | A0A8I6Z7D7 | Peptidase A1 domain-containing protein | 1.40 | 4.96 | 2.53E-05 |
| 1HG0080920 | A0A8I6WEJ1 | WRKY domain-containing protein | 1.39 | 4.94 | 2.85E-05 |
| 7HG0664270 | A0A8I6Z0S5 | NADH:flavin oxidoreductase/NADH oxidase N-terminal domain-containing protein | 1.39 | 4.21 | 2.72E-04 |
| 5HG0484690 | F2D024 | Predicted protein | 1.39 | 7.32 | 6.34E-05 |
| 6HG0543180 | A0A8I7BAW1 | Germin-like protein | 1.39 | 2.45 | 4.31E-02 |
| 1HG0004730 | A0A8I6WNW4 | RING-type E3 ubiquitin transferase (EC 2.3.2.27) | 1.38 | 6.95 | 3.69E-05 |
| 6HG0574110 | A0A8I6YRX4 | U-box domain-containing protein (EC 2.3.2.27) (RING-type E3 ubiquitin transferase PUB) | 1.38 | 3.66 | 2.17E-03 |
| 6HG0615560 | F2DMM7 | Predicted protein | 1.38 | 5.61 | 1.27E-05 |
| 5HG0511770 | F2D0E7 | UDP-glucose 6-dehydrogenase (EC 1.1.1.22) | 1.38 | 8.61 | 4.69E-04 |
| 2HG0191460 | A0A8I6X6R0 | Cationic amino acid transporter C-terminal domain-containing protein | 1.38 | 2.89 | 1.90E-02 |
| 2HG0124450 | F2DAN9 | Predicted protein | 1.38 | 4.31 | 2.27E-04 |
| 7HG0733220 | A0A8I6YRF0 | Ubiquitin-like domain-containing protein | 1.38 | 4.40 | 1.68E-04 |
| 5HG0464590 | M0W4N1 | Uncharacterized protein | 1.37 | 2.93 | 1.81E-02 |
| 7HG0663990 | A0A8I6YD14 | Scarecrow-like protein 9 | 1.37 | 4.98 | 3.69E-05 |
| 5HG0479040 | F2DA97 | Predicted protein | 1.36 | 5.87 | 1.70E-05 |
| 4HG0409490 | A0A8I6XIB9 | NAC domain-containing protein | 1.36 | 4.76 | 6.93E-05 |
| 5HG0517040 | Q6X9V9 | Predicted protein (Syntaxin) | 1.36 | 6.02 | 1.82E-05 |
| 3HG0267290 | F2DXR6 | Predicted protein | 1.36 | 5.91 | 1.85E-05 |
| 3HG0308630 | A0A8I6X5Z2 | Auxin response factor | 1.36 | 3.93 | 1.11E-03 |
| 4HG0400780 | F2E8Z2 | Predicted protein | 1.36 | 6.18 | 2.16E-05 |
| 3HG0240640 | A0A8I6WQ24 | Glycosyltransferase (EC 2.4.1.-) | 1.35 | 4.96 | 4.90E-05 |
| 1HG0018260 | A0A8I6WN09 | Uncharacterized protein | 1.35 | 6.62 | 3.91E-05 |
| 6HG0607680 | A0A8I6Y491 | SANT domain-containing protein | 1.34 | 3.20 | 1.21E-02 |
| 3HG0280930 | A0A8I6X2H9 | Uncharacterized protein | 1.34 | 4.15 | 6.28E-04 |
| 6HG0632070 | A0A8I6YXC9 | Protein kinase domain-containing protein | 1.34 | 6.01 | 2.83E-05 |
| 6HG0574250 | A0A8I6Y0W0 | Uncharacterized protein | 1.33 | 5.07 | 5.69E-05 |
| 2HG0098030 | A0A8I6WMK6 | Rx N-terminal domain-containing protein | 1.33 | 2.80 | 3.12E-02 |
| 5HG0421580 | A0A8I6Y5M0 | DUF1618 domain-containing protein | 1.33 | 4.39 | 3.46E-04 |
| 7HG0665380 | A0A8I6YAK7 | AP2/ERF domain-containing protein | 1.32 | 5.61 | 3.44E-05 |
| 2HG0186080 | A0A8I6XAN5 | Fe2OG dioxygenase domain-containing protein | 1.32 | 3.40 | 8.73E-03 |
| 6HG0542640 | A0A8I6Y973 | Uncharacterized protein | 1.31 | 5.44 | 4.76E-05 |
| 2HG0197410 | A0A8I6WI79 | non-specific serine/threonine protein kinase (EC 2.7.11.1) | 1.31 | 2.70 | 4.31E-02 |
| 1HG0061300 | M0YZK1 | Uncharacterized protein | 1.31 | 2.75 | 3.98E-02 |
| 2HG0182010 | A0A8I6WQT6 | Phenylalanine ammonia-lyase (EC 4.3.1.24) | 1.30 | 5.25 | 6.93E-05 |
| 3HG0294340 | A0A8I6XGK9 | Peroxidase (EC 1.11.1.7) | 1.30 | 3.96 | 1.95E-03 |
| 3HG0319710 | A0A8I6XJE8 | glutathione transferase (EC 2.5.1.18) | 1.30 | 3.44 | 9.34E-03 |
| 5HG0433130 | A0A8I6XNQ1 | EF-hand domain-containing protein | 1.29 | 3.73 | 4.15E-03 |
| 4HG0338560 | A0A8I6XF27 | Uncharacterized protein | 1.29 | 2.98 | 2.80E-02 |
| 5HG0520830 | A0A8I6XLY1 | DUF569 domain-containing protein | 1.28 | 4.79 | 2.11E-04 |
| 2HG0213040 | A0A8I6XM05 | AP2/ERF domain-containing protein | 1.28 | 6.08 | 6.93E-05 |
| 5HG0460690 | A0A8I6XAW6 | PGG domain-containing protein | 1.28 | 3.04 | 2.74E-02 |
| 2HG0122480 | A0A8I6XAZ7 | Glycosyltransferase (EC 2.4.1.-) | 1.28 | 3.91 | 2.89E-03 |
| 3HG0233040 | A0A8I6X4I0 | RING-type E3 ubiquitin transferase (EC 2.3.2.27) | 1.28 | 4.15 | 1.40E-03 |
| 1HG0092830 | A0A287GRI5 | Glycosyltransferase (EC 2.4.1.-) | 1.27 | 6.15 | 8.91E-05 |
| 2HG0101890 | F2DT93 | Predicted protein | 1.27 | 5.14 | 1.31E-04 |
| 3HG0229420 | F2DE42 | Predicted protein | 1.27 | 5.70 | 8.17E-05 |
| 2HG0125660 | A0A8I7B2F3 | DUF1677 family protein | 1.27 | 4.57 | 4.66E-04 |
| 5HG0448660 | A0A8I6XM18 | Calmodulin-binding domain-containing protein | 1.27 | 3.19 | 2.11E-02 |
| 6HG0540160 | A0A8I6YFJ2 | Rhodanese domain-containing protein | 1.27 | 4.88 | 2.21E-04 |
| 7HG0715110 | A0A8I6Z494 | Double-strand break repair protein | 1.27 | 3.37 | 1.42E-02 |
| 2HG0138730 | A0A8I6WC76 | Uncharacterized protein | 1.26 | 2.97 | 3.39E-02 |
| 5HG0481730 | F2DN89 | Glutamate receptor | 1.25 | 4.21 | 1.48E-03 |
| 3HG0297360 | F2CVH1 | Predicted protein | 1.25 | 5.20 | 1.53E-04 |
| 7HG0647190 | A0A287VK78 | phosphopyruvate hydratase (EC 4.2.1.11) | 1.25 | 5.48 | 1.14E-04 |
| 7HG0682090 | A0A8I6Z9S6 | Oxysterol-binding protein | 1.25 | 5.52 | 1.14E-04 |
| 1HG0029480 | A0A8I7B2R7 | DDE Tnp4 domain-containing protein | 1.25 | 5.28 | 1.53E-04 |
| 2HG0104870 | A0A8I6WK84 | cinnamyl-alcohol dehydrogenase (EC 1.1.1.195) | 1.25 | 3.31 | 1.86E-02 |
| 1HG0051180 | F2D2G5 | RHOMBOID-like protein (EC 3.4.21.105) | 1.24 | 2.92 | 4.32E-02 |
| 2HG0211040 | F2DRN3 | Predicted protein | 1.24 | 4.51 | 7.87E-04 |
| 1HG0055890 | F2E6C0 | Proline dehydrogenase (EC 1.5.5.2) | 1.23 | 8.67 | 2.92E-03 |
| 7HG0709440 | A0A8I6YQ87 | RING-type E3 ubiquitin transferase (EC 2.3.2.27) | 1.23 | 3.91 | 4.80E-03 |
| 6HG0622920 | A0A8I6YG31 | Trehalose 6-phosphate phosphatase | 1.22 | 9.99 | 1.03E-02 |
| 1HG0076630 | A0A8I6W7G6 | Protein kinase domain-containing protein | 1.22 | 3.17 | 3.11E-02 |
| 1HG0089010 | A0A8I7B464 | Uncharacterized protein | 1.22 | 3.22 | 2.81E-02 |
| 4HG0392580 | A0A8I6XDQ5 | Thaumatin-like protein | 1.22 | 3.16 | 3.23E-02 |
| 7HG0662420 | F2E513 | Predicted protein | 1.21 | 5.10 | 3.38E-04 |
| 7HG0665050 | F2E7Z5 | Germin-like protein | 1.21 | 4.49 | 1.25E-03 |
| 1HG0056490 | A0A8I6WJ59 | Glucuronosyltransferase PGSIP8 | 1.20 | 4.46 | 1.48E-03 |
| 7HG0648120 | A0A8I6YZ57 | glutathione transferase (EC 2.5.1.18) | 1.20 | 4.30 | 2.24E-03 |
| 2HG0183740 | A0A8I6WG70 | Major facilitator superfamily (MFS) profile domain-containing protein | 1.20 | 3.19 | 3.44E-02 |
| 4HG0384700 | A0A8I7B7B1 | protein-serine/threonine phosphatase (EC 3.1.3.16) | 1.20 | 4.75 | 7.98E-04 |
| 3HG0240580 | D3WYW1 | Glycosyltransferase (EC 2.4.1.-) | 1.19 | 4.80 | 7.58E-04 |
| 1HG0015720 | F2CR53 | Predicted protein | 1.19 | 5.97 | 2.86E-04 |
| 4HG0413910 | F2CZP9 | beta-fructofuranosidase (EC 3.2.1.26) | 1.19 | 7.40 | 1.21E-03 |
| 5HG0514220 | A0A8I7B9Y0 | F-box domain-containing protein | 1.18 | 3.65 | 1.41E-02 |
| 5HG0463010 | F2EEW5 | Predicted protein | 1.18 | 3.28 | 3.12E-02 |
| 3HG0230870 | F2D304 | Predicted protein | 1.18 | 6.59 | 5.38E-04 |
| 2HG0208700 | F2EIZ1 | Predicted protein | 1.18 | 6.46 | 5.03E-04 |
| 4HG0400000 | A0A8I6XEH6 | DOMON domain-containing protein | 1.17 | 4.80 | 9.49E-04 |
| 3HG0247120 | F2E516 | Predicted protein | 1.17 | 4.21 | 3.78E-03 |
| 3HG0245440 | F2E1F6 | Predicted protein | 1.17 | 6.51 | 5.58E-04 |
| 1HG0038590 | A0A8I7B0T2 | HMA domain-containing protein | 1.17 | 8.35 | 4.15E-03 |
| 3HG0299690 | F2D7J1 | Predicted protein | 1.17 | 4.09 | 5.37E-03 |
| 7HG0731730 | A0A8I6ZE90 | Bifunctional inhibitor/plant lipid transfer protein/seed storage helical domain-containing protein | 1.17 | 7.57 | 1.95E-03 |
| 4HG0390960 | A0A8I6Y0N3 | PGG domain-containing protein | 1.17 | 5.50 | 4.62E-04 |
| 2HG0099640 | A0A8I6X0J4 | Glycosyltransferase (EC 2.4.1.-) | 1.16 | 4.68 | 1.46E-03 |
| 6HG0609420 | F2D579 | Predicted protein | 1.16 | 4.14 | 5.02E-03 |
| 4HG0395160 | A0A8I6XDX4 | Protein kinase domain-containing protein | 1.16 | 5.66 | 4.68E-04 |
| 5HG0521630 | A0A8I6YL71 | Galectin domain-containing protein | 1.16 | 3.45 | 2.74E-02 |
| 7HG0656670 | F2CPZ7 | galactinol--sucrose galactosyltransferase (EC 2.4.1.82) | 1.15 | 7.21 | 1.61E-03 |
| 7HG0707770 | A0A8I7BEQ5 | Uncharacterized protein | 1.15 | 3.35 | 3.44E-02 |
| 2HG0215200 | A0A8I6X145 | Uncharacterized protein | 1.15 | 5.24 | 7.38E-04 |
| 2HG0212850 | A0A8I6XFN6 | glutathione transferase (EC 2.5.1.18) | 1.15 | 4.09 | 7.01E-03 |
| 4HG0337000 | A0A8I7BBJ6 | DUF292 domain-containing protein | 1.14 | 7.11 | 1.55E-03 |
| 6HG0553820 | A0A8I6XYT5 | Leucine-rich repeat-containing N-terminal plant-type domain-containing protein | 1.14 | 7.37 | 2.22E-03 |
| 3HG0226060 | A0A8I6WN69 | EF-hand domain-containing protein | 1.14 | 5.24 | 8.34E-04 |
| 7HG0718770 | A0A287X8U4 | Uncharacterized protein | 1.14 | 4.05 | 8.51E-03 |
| 3HG0229410 | M0XS96 | OB domain-containing protein | 1.13 | 5.73 | 6.76E-04 |
| 7HG0746630 | A0A8I6YUY9 | Peptidase A1 domain-containing protein | 1.13 | 5.39 | 7.95E-04 |
| 5HG0448210 | A0A8I7BDZ6 | Heptahelical transmembrane protein 4 | 1.13 | 4.66 | 2.38E-03 |
| 2HG0167390 | F2EKH1 | Predicted protein | 1.13 | 3.63 | 2.35E-02 |
| 4HG0394970 | M0ZBM5 | Allene oxide synthase | 1.12 | 5.73 | 7.86E-04 |
| 2HG0186310 | F2E442 | Predicted protein | 1.12 | 5.84 | 8.00E-04 |
| 5HG0463440 | F2E4T2 | Predicted protein | 1.12 | 3.85 | 1.49E-02 |
| 3HG0322360 | A0A8I6XUF5 | Uncharacterized protein | 1.12 | 3.50 | 3.13E-02 |
| 3HG0233030 | A0A8I6WPA8 | RING-type E3 ubiquitin transferase (EC 2.3.2.27) | 1.12 | 4.17 | 7.41E-03 |
| 1HG0073940 | F2DZX6 | Predicted protein | 1.12 | 7.86 | 4.80E-03 |
| 3HG0287850 | A0A287LJS0 | Uncharacterized protein | 1.12 | 6.37 | 1.11E-03 |
| 5HG0504160 | F2CUP3 | Histone H4 | 1.12 | 3.78 | 1.83E-02 |
| 6HG0618650 | A0A8I6Y6Q6 | aminocyclopropanecarboxylate oxidase (EC 1.14.17.4) (Ethylene-forming enzyme) | 1.11 | 4.69 | 2.72E-03 |
| 1HG0074300 | A0A8I6WSK6 | Uncharacterized protein | 1.10 | 3.63 | 2.80E-02 |
| 2HG0177740 | A0A8I6XF63 | Uncharacterized protein | 1.10 | 5.85 | 1.05E-03 |
| 5HG0525180 | A0A8I6XUD3 | Protein LURP-one-related 15 | 1.10 | 5.28 | 1.34E-03 |
| 7HG0668040 | F2DER0 | Predicted protein | 1.10 | 4.13 | 9.80E-03 |
| 5HG0440150 | F2CPQ5 | Predicted protein | 1.10 | 3.77 | 2.11E-02 |
| 2HG0186540 | A0A8I6XFZ4 | Uncharacterized protein | 1.10 | 4.66 | 3.33E-03 |
| 5HG0480600 | A0A8I6XNE8 | Protein TIFY (Jasmonate ZIM domain-containing protein) | 1.10 | 3.43 | 4.29E-02 |
| 3HG0236310 | F2CUW9 | Predicted protein | 1.10 | 4.59 | 3.84E-03 |
| 2HG0208650 | A0A8I6XRJ9 | HMA domain-containing protein | 1.09 | 4.73 | 3.12E-03 |
| 1HG0027490 | F2DMD5 | Predicted protein | 1.09 | 3.88 | 1.83E-02 |
| 4HG0397090 | F2E8E4 | Predicted protein | 1.09 | 3.98 | 1.48E-02 |
| 5HG0493870 | A0A8I6YAG0 | Auxin efflux carrier component | 1.09 | 3.43 | 4.57E-02 |
| 7HG0669750 | A0A8I6YKZ8 | Fungal lipase-like domain-containing protein | 1.09 | 4.90 | 2.66E-03 |
| 4HG0334010 | A0A8I6XEK2 | Uncharacterized protein | 1.09 | 5.54 | 1.42E-03 |
| 6HG0574410 | F2E7K4 | Predicted protein | 1.08 | 6.24 | 1.58E-03 |
| 1HG0019970 | A0A8I6WRE3 | Uncharacterized protein | 1.08 | 4.27 | 8.91E-03 |
| 1HG0077190 | A0A287GAZ3 | alpha,alpha-trehalose-phosphate synthase (UDP-forming) (EC 2.4.1.15) | 1.08 | 7.45 | 4.80E-03 |
| 6HG0574360 | F2E2C9 | Predicted protein | 1.08 | 5.40 | 1.61E-03 |
| 4HG0343130 | F2EBH8 | Predicted protein | 1.08 | 4.32 | 8.23E-03 |
| 4HG0383900 | A0A8I6YHC2 | Uncharacterized protein | 1.08 | 6.05 | 1.52E-03 |
| 4HG0412490 | A0A8I6YAX5 | Fe2OG dioxygenase domain-containing protein | 1.08 | 7.94 | 7.71E-03 |
| 2HG0126450 | A0A8I6XE59 | Sulfotransferase (EC 2.8.2.-) | 1.08 | 3.50 | 4.35E-02 |
| 3HG0298790 | A0A8I6WUH4 | amidophosphoribosyltransferase (EC 2.4.2.14) | 1.07 | 6.80 | 2.84E-03 |
| 4HG0396130 | A0A8I6Y8R2 | Major facilitator superfamily (MFS) profile domain-containing protein | 1.07 | 8.07 | 9.34E-03 |
| 1HG0064880 | A0A8I6WK72 | Alpha/beta hydrolase fold-3 domain-containing protein | 1.07 | 4.84 | 3.39E-03 |
| 1HG0051910 | A0A8I6WII2 | glutathione transferase (EC 2.5.1.18) | 1.07 | 6.02 | 1.75E-03 |
| 5HG0510240 | A0A8I6Y671 | Dirigent protein | 1.06 | 4.25 | 1.14E-02 |
| 7HG0664700 | A0A8I6YAI3 | RING-type E3 ubiquitin transferase (EC 2.3.2.27) | 1.06 | 5.59 | 2.01E-03 |
| 2HG0110850 | F2DU39 | Predicted protein | 1.06 | 5.41 | 2.22E-03 |
| 3HG0302620 | A0A287M2R6 | Uncharacterized protein | 1.06 | 5.40 | 2.24E-03 |
| 2HG0208660 | F2D8L6 | Predicted protein | 1.06 | 7.42 | 6.41E-03 |
| 2HG0182020 | A0A287IVK8 | Phenylalanine ammonia-lyase (EC 4.3.1.24) | 1.06 | 4.39 | 9.34E-03 |
| 7HG0683070 | F2DWN8 | Predicted protein | 1.06 | 6.61 | 3.03E-03 |
| 5HG0536910 | A0A8I6Y552 | Cytokinin riboside 5'-monophosphate phosphoribohydrolase (EC 3.2.2.n1) | 1.05 | 5.96 | 2.17E-03 |
| 3HG0304380 | A0A8I7B663 | WRKY domain-containing protein | 1.05 | 8.63 | 1.73E-02 |
| 2HG0127020 | F2CZ17 | Predicted protein | 1.05 | 3.58 | 4.56E-02 |
| 7HG0635380 | M0X3V0 | Beta-fructofuranosidase | 1.05 | 4.18 | 1.47E-02 |
| 6HG0628070 | A0A8I6YZ25 | Uncharacterized protein | 1.04 | 5.14 | 3.34E-03 |
| 5HG0485780 | F2D6S5 | aminocyclopropanecarboxylate oxidase (EC 1.14.17.4) (Ethylene-forming enzyme) | 1.04 | 3.56 | 4.93E-02 |
| 4HG0396540 | M0YH92 | Sugar phosphate transporter domain-containing protein | 1.04 | 5.27 | 3.10E-03 |
| 3HG0219640 | F2E750 | Predicted protein | 1.03 | 3.75 | 3.75E-02 |
| 6HG0602910 | A0A8I6XVN1 | Uncharacterized protein | 1.03 | 6.38 | 3.52E-03 |
| 1HG0051870 | A0A8I7B355 | glutathione transferase (EC 2.5.1.18) | 1.03 | 6.22 | 3.39E-03 |
| 5HG0522100 | A0A8I6YWA7 | Germin-like protein | 1.03 | 7.19 | 7.30E-03 |
| 2HG0128980 | F2DJ70 | Predicted protein | 1.03 | 4.29 | 1.47E-02 |
| 2HG0171680 | A0A8I6X4L7 | Lipoxygenase (EC 1.13.11.-) | 1.02 | 6.70 | 4.93E-03 |
| 6HG0617010 | F2EGH1 | Predicted protein | 1.02 | 5.06 | 5.11E-03 |
| 1HG0047110 | A0A8I6WFV9 | AB hydrolase-1 domain-containing protein | 1.02 | 5.80 | 3.46E-03 |
| 3HG0266950 | A0A287KZR6 | Calmodulin-binding protein | 1.02 | 7.21 | 8.40E-03 |
| 2HG0109340 | A0A8I6WR00 | Wound-induced protein 1 | 1.02 | 6.04 | 3.66E-03 |
| 5HG0490260 | A0A8I6XTT5 | FAD-binding PCMH-type domain-containing protein | 1.02 | 4.45 | 1.25E-02 |
| 4HG0412340 | F2DP63 | Predicted protein | 1.01 | 6.70 | 5.50E-03 |
| 5HG0518070 | F2CXS3 | Glycosyltransferase (EC 2.4.1.-) | 1.01 | 4.18 | 2.00E-02 |
| 6HG0548800 | A0A8I6YPN3 | Uncharacterized protein | 1.01 | 3.99 | 2.90E-02 |
| 1HG0089440 | F2CQB5 | Predicted protein | 1.01 | 5.82 | 3.78E-03 |
| 1HG0091980 | A0A8I6XAB7 | Myb-like domain-containing protein | 1.01 | 5.81 | 3.83E-03 |
| 2HG0177660 | A0A8I6WVR0 | EF-hand domain-containing protein | 1.01 | 4.86 | 7.41E-03 |
| 1HG0084760 | A0A8I6WIW3 | Uncharacterized protein | 1.01 | 4.03 | 2.80E-02 |
| 4HG0415460 | A0A8I6XLL8 | Protein kinase domain-containing protein | 1.00 | 5.41 | 4.53E-03 |
| 6HG0547990 | A0A8I6YYY1 | RING-type E3 ubiquitin transferase (EC 2.3.2.27) | 1.00 | 4.21 | 2.06E-02 |
| 2HG0121540 | A0A8I6WPM9 | Uncharacterized protein | 1.00 | 3.79 | 4.44E-02 |
| 7HG0750360 | M0YGH3 | 15-cis-phytoene synthase (EC 2.5.1.32) | -1.00 | 4.53 | 1.24E-02 |
| 3HG0261980 | F2E119 | Serine/threonine-protein phosphatase (EC 3.1.3.16) | -1.01 | 4.17 | 2.11E-02 |
| 5HG0487460 | A0A8I6XGE8 | Ternary complex factor MIP1 leucine-zipper domain-containing protein | -1.01 | 4.81 | 7.44E-03 |
| 2HG0182920 | M0VNP8 | Large ribosomal RNA subunit accumulation protein YCED homolog 2, chloroplastic | -1.01 | 3.95 | 3.07E-02 |
| 2HG0206550 | A0A8I6XEI4 | shikimate kinase (EC 2.7.1.71) | -1.01 | 3.94 | 3.11E-02 |
| 5HG0440860 | M0V9E2 | Kinesin-like protein | -1.02 | 5.49 | 3.76E-03 |
| 1HG0050670 | M0V3I5 | Uncharacterized protein | -1.02 | 4.09 | 2.10E-02 |
| 5HG0515030 | F2CXL1 | Obg-like ATPase 1 | -1.03 | 3.74 | 3.99E-02 |
| 4HG0377630 | A0A8I6XCC5 | Uncharacterized protein | -1.03 | 3.64 | 4.70E-02 |
| 1HG0073870 | A0A8I6WKU9 | Delta-1-pyrroline-5-carboxylate synthase [Includes: Glutamate 5-kinase (GK) (EC 2.7.2.11) (Gamma-glutamyl kinase); Gamma-glutamyl phosphate reductase (GPR) (EC 1.2.1.41) (Glutamate-5-semialdehyde dehydrogenase) (Glutamyl-gamma-semialdehyde dehydrogenase)] | -1.03 | 5.57 | 2.93E-03 |
| 7HG0740500 | A0A8I6Z6R4 | F-box domain-containing protein | -1.03 | 4.94 | 4.91E-03 |
| 2HG0202350 | A0A8I6XJ34 | Glutamine synthetase (EC 6.3.1.2) | -1.03 | 9.19 | 2.87E-02 |
| 5HG0483990 | A0A8I6XFY8 | non-specific serine/threonine protein kinase (EC 2.7.11.1) | -1.03 | 4.85 | 5.31E-03 |
| 2HG0161380 | A0A8I6WUL0 | VDE lipocalin domain-containing protein | -1.04 | 5.49 | 2.92E-03 |
| 5HG0443280 | A0A8I7B8E6 | PsbP C-terminal domain-containing protein | -1.04 | 4.47 | 9.17E-03 |
| 5HG0491450 | A0A8I6XPV2 | NAD(P)-binding domain-containing protein | -1.04 | 4.45 | 9.37E-03 |
| 4HG0376960 | A0A8I6YGK0 | RRM domain-containing protein | -1.04 | 4.41 | 1.01E-02 |
| 3HG0227400 | F2CSG7 | Non-specific lipid-transfer protein | -1.04 | 7.21 | 6.00E-03 |
| 3HG0302060 | A0A8I6X9N2 | Delta-1-pyrroline-5-carboxylate synthase [Includes: Glutamate 5-kinase (GK) (EC 2.7.2.11) (Gamma-glutamyl kinase); Gamma-glutamyl phosphate reductase (GPR) (EC 1.2.1.41) (Glutamate-5-semialdehyde dehydrogenase) (Glutamyl-gamma-semialdehyde dehydrogenase)] | -1.05 | 3.84 | 2.73E-02 |
| 5HG0502720 | M0Y4N4 | Cytokinin riboside 5'-monophosphate phosphoribohydrolase (EC 3.2.2.n1) | -1.06 | 4.08 | 1.59E-02 |
| 7HG0665750 | A0A287W3F1 | Uncharacterized protein | -1.06 | 5.28 | 2.39E-03 |
| 2HG0102500 | A0A8I6W9H5 | RuBisCO large subunit-binding protein subunit alpha, chloroplastic | -1.08 | 6.17 | 1.51E-03 |
| 3HG0318900 | A0A287MJH1 | Glucan endo-1,3-beta-D-glucosidase | -1.09 | 3.38 | 4.93E-02 |
| 5HG0425790 | A0A8I6Y4Z4 | DUF6598 domain-containing protein | -1.09 | 3.61 | 3.19E-02 |
| 5HG0482670 | F2CSG2 | Cellulose synthase (EC 2.4.1.12) | -1.09 | 4.94 | 2.32E-03 |
| 4HG0333190 | A0A8I6XED1 | NusB/RsmB/TIM44 domain-containing protein | -1.09 | 4.99 | 2.17E-03 |
| 5HG0471050 | F2DD45 | Predicted protein | -1.09 | 5.56 | 1.27E-03 |
| 4HG0380970 | A0A8I6XK45 | Hexosyltransferase (EC 2.4.1.-) | -1.10 | 3.94 | 1.53E-02 |
| 4HG0412610 | A0A8I6XIM1 | NAC-A/B domain-containing protein | -1.10 | 5.27 | 1.38E-03 |
| 5HG0432540 | F2CW52 | Predicted protein | -1.11 | 3.34 | 4.74E-02 |
| 1HG0092170 | A0A8I6WYN4 | rRNA N-glycosidase | -1.11 | 3.63 | 2.62E-02 |
| 2HG0193390 | F2CY55 | Predicted protein | -1.11 | 5.82 | 9.42E-04 |
| 5HG0513900 | A0A8I7BFF8 | S-adenosyl-L-methionine-dependent methyltransferase | -1.12 | 5.46 | 9.61E-04 |
| 1HG0066180 | A0A8I6WDL1 | HMA domain-containing protein | -1.12 | 4.81 | 2.01E-03 |
| 7HG0636480 | A0A8I7BHZ6 | Leucine-rich repeat-containing N-terminal plant-type domain-containing protein | -1.13 | 6.15 | 8.34E-04 |
| 3HG0288960 | F2DMG1 | Predicted protein | -1.14 | 5.03 | 1.11E-03 |
| 2HG0190670 | A0A8I7B7Z2 | DUF642 domain-containing protein | -1.14 | 3.77 | 1.52E-02 |
| 2HG0209260 | F2EFN1 | Predicted protein | -1.14 | 4.50 | 2.69E-03 |
| 6HG0574460 | F2D426 | Glutaredoxin-dependent peroxiredoxin (EC 1.11.1.25) | -1.15 | 5.33 | 6.25E-04 |
| 7HG0652120 | A0A8I6Z728 | 2Fe-2S ferredoxin-type domain-containing protein | -1.16 | 6.93 | 1.03E-03 |
| 2HG0192050 | A0A8I6WX09 | Uncharacterized protein | -1.16 | 3.42 | 2.73E-02 |
| 3HG0277690 | A0A8I7BA40 | 1,4-dihydroxy-2-naphthoyl-CoA synthase, peroxisomal | -1.16 | 4.10 | 5.46E-03 |
| 1HG0080860 | A0A8I6WLX1 | Sulfotransferase (EC 2.8.2.-) | -1.17 | 3.61 | 1.78E-02 |
| 4HG0354250 | A0A8I6YF25 | Protein kinase domain-containing protein | -1.17 | 3.28 | 3.44E-02 |
| 5HG0469020 | F2CT91 | Ribulose bisphosphate carboxylase small subunit, chloroplastic (RuBisCO small subunit) | -1.18 | 11.46 | 3.29E-02 |
| 4HG0340410 | A0A287N786 | rRNA methyltransferase | -1.18 | 3.65 | 1.46E-02 |
| 5HG0527630 | A0A8I6XMJ8 | Protein kinase domain-containing protein | -1.19 | 4.84 | 7.18E-04 |
| 3HG0274780 | A0A287L5I7 | Carbonic anhydrase (EC 4.2.1.1) (Carbonate dehydratase) | -1.19 | 10.82 | 2.10E-02 |
| 5HG0448480 | F2CVF0 | Predicted protein | -1.20 | 3.06 | 4.31E-02 |
| 7HG0713590 | A0A8I6YQN4 | Enoyl reductase (ER) domain-containing protein | -1.21 | 5.00 | 3.91E-04 |
| 3HG0270730 | A0A8I6XMN2 | Pentatricopeptide repeat-containing protein | -1.21 | 3.34 | 2.19E-02 |
| 7HG0748080 | A0A8I6YI05 | HMA domain-containing protein | -1.22 | 4.59 | 8.65E-04 |
| 2HG0176940 | A0A8I6X558 | Malate synthase (EC 2.3.3.9) | -1.22 | 7.51 | 9.46E-04 |
| 1HG0027000 | M0VUM7 | Uncharacterized protein | -1.23 | 3.16 | 2.99E-02 |
| 5HG0520260 | F2E5C6 | Predicted protein | -1.23 | 3.93 | 4.16E-03 |
| 2HG0192890 | A0A8I6XGH9 | Xyloglucan endotransglucosylase/hydrolase (EC 2.4.1.207) | -1.23 | 3.96 | 3.78E-03 |
| 4HG0405370 | A0A8I6Y254 | Pentatricopeptide repeat-containing protein | -1.24 | 3.95 | 3.74E-03 |
| 6HG0541050 | F2E960 | Predicted protein | -1.24 | 3.96 | 3.55E-03 |
| 4HG0405920 | A0A8I6XKJ3 | Hexosyltransferase (EC 2.4.1.-) | -1.26 | 3.52 | 1.03E-02 |
| 7HG0662060 | A0A8I6Z802 | Pentacotripeptide-repeat region of PRORP domain-containing protein | -1.26 | 3.97 | 2.84E-03 |
| 1HG0082230 | A0A8I7B3Z7 | DC-UbP/UBTD2 N-terminal domain-containing protein | -1.26 | 3.02 | 3.05E-02 |
| 3HG0297000 | A0A8I6X1S6 | Protein LURP-one-related 8 | -1.27 | 3.02 | 2.92E-02 |
| 7HG0649250 | F2EIK6 | Superoxide dismutase (EC 1.15.1.1) | -1.28 | 3.43 | 1.10E-02 |
| 1HG0059680 | M0W8S9 | eRF1 domain-containing protein | -1.28 | 4.22 | 1.03E-03 |
| 2HG0197190 | A0A8I6XCR9 | RRM domain-containing protein | -1.28 | 4.49 | 4.62E-04 |
| 7HG0653480 | A0A8I6YJ64 | Short-chain dehydrogenase/reductase (EC 1.1.1.-) | -1.29 | 2.74 | 4.38E-02 |
| 2HG0139380 | A0A8I6XFM0 | Isocitrate lyase | -1.29 | 3.72 | 4.36E-03 |
| 7HG0642890 | A0A287VHY2 | Acid phosphatase | -1.30 | 3.83 | 3.00E-03 |
| 5HG0446040 | A0A8I7B8H0 | COBRA-like protein | -1.30 | 3.35 | 1.17E-02 |
| 1HG0059670 | I3VIX5 | cytokinin dehydrogenase (EC 1.5.99.12) | -1.31 | 3.20 | 1.56E-02 |
| 3HG0253860 | F2EAV7 | Predicted protein | -1.31 | 3.84 | 2.46E-03 |
| 1HG0040570 | A0A8I6WGY7 | SAP domain-containing protein | -1.32 | 3.84 | 2.29E-03 |
| 4HG0411020 | F2CZZ9 | Predicted protein | -1.32 | 3.13 | 1.70E-02 |
| 5HG0473680 | A0A8I6XEF4 | protein-serine/threonine phosphatase (EC 3.1.3.16) | -1.32 | 6.01 | 3.53E-05 |
| 3HG0250140 | A0A8I6X694 | NmrA-like domain-containing protein | -1.32 | 4.57 | 2.02E-04 |
| 5HG0464380 | A0A8I6XBB4 | Chalcone-flavonone isomerase family protein | -1.34 | 4.04 | 9.42E-04 |
| 5HG0436160 | A0A8I6XIC5 | Uncharacterized protein | -1.34 | 3.62 | 3.68E-03 |
| 4HG0395580 | A0A8I6X3P9 | Protein kinase domain-containing protein | -1.34 | 3.40 | 7.00E-03 |
| 1HG0024740 | F2D5Y4 | Predicted protein | -1.35 | 2.79 | 2.76E-02 |
| 7HG0712010 | M0YG85 | DJ-1/PfpI domain-containing protein | -1.35 | 4.53 | 1.53E-04 |
| 6HG0623560 | A0A8I6XYF6 | Clu domain-containing protein | -1.36 | 6.23 | 2.16E-05 |
| 6HG0621560 | A0A8I6YFX8 | Glycosyltransferase (EC 2.4.1.-) | -1.36 | 3.00 | 1.69E-02 |
| 2HG0112150 | F2DL08 | Alpha-galactosidase (EC 3.2.1.22) (Melibiase) | -1.37 | 2.91 | 1.98E-02 |
| 7HG0705460 | F2D734 | Predicted protein | -1.37 | 6.52 | 2.60E-05 |
| 6HG0542030 | A0A8I6Y660 | Beta-fructofuranosidase | -1.37 | 3.19 | 1.02E-02 |
| 5HG0499440 | F2DDX8 | Endoglucanase (EC 3.2.1.4) | -1.37 | 2.93 | 1.83E-02 |
| 6HG0633810 | F2DRP6 | Sucrose synthase (EC 2.4.1.13) | -1.37 | 2.60 | 3.47E-02 |
| 3HG0257380 | A0A8I6XMB7 | Uncharacterized protein | -1.38 | 2.41 | 4.62E-02 |
| 7HG0686900 | A0A8I6YCP1 | Ubiquitin-like domain-containing protein | -1.39 | 5.14 | 2.00E-05 |
| 3HG0244080 | A0A287KIB4 | Uncharacterized protein | -1.39 | 4.51 | 9.93E-05 |
| 7HG0637280 | A0A287VE25 | Alpha-dioxygenase 1 | -1.40 | 4.25 | 2.05E-04 |
| 5HG0433990 | F2EE60 | alanine transaminase (EC 2.6.1.2) | -1.41 | 4.27 | 1.67E-04 |
| 6HG0590910 | A0A8I7BBX7 | NPH3 domain-containing protein | -1.41 | 5.04 | 1.63E-05 |
| 3HG0227360 | F2DCX3 | Non-specific lipid-transfer protein | -1.41 | 6.85 | 1.96E-05 |
| 2HG0138340 | F2E4J5 | Predicted protein | -1.42 | 2.70 | 2.11E-02 |
| 2HG0165730 | A0A8I6X434 | RAB6-interacting golgin | -1.43 | 2.77 | 1.75E-02 |
| 5HG0494890 | A0A8I6Y428 | Serine aminopeptidase S33 domain-containing protein | -1.44 | 4.36 | 7.53E-05 |
| 3HG0224210 | A0A8I6XA81 | Peroxisomal membrane protein PEX14 (Peroxin-14) | -1.44 | 2.41 | 3.32E-02 |
| 1HG0090670 | F2DJX7 | protein-serine/threonine phosphatase (EC 3.1.3.16) | -1.45 | 2.25 | 4.18E-02 |
| 2HG0172300 | A0A8I6WVA9 | Uncharacterized protein | -1.45 | 4.04 | 2.27E-04 |
| 7HG0659160 | A0A8I6Y413 | NAD-dependent epimerase/dehydratase domain-containing protein | -1.46 | 2.33 | 3.49E-02 |
| 1HG0037500 | F2D629 | Ribulose bisphosphate carboxylase small subunit, chloroplastic (RuBisCO small subunit) | -1.46 | 8.06 | 6.93E-05 |
| 3HG0219130 | A0A287JVY4 | Peptidyl-prolyl cis-trans isomerase (PPIase) (EC 5.2.1.8) | -1.47 | 3.69 | 7.55E-04 |
| 6HG0614510 | A0A8I6YM48 | peptidylprolyl isomerase (EC 5.2.1.8) | -1.47 | 5.17 | 4.31E-06 |
| 1HG0041280 | D9IXC7 | Cellulose synthase (EC 2.4.1.12) | -1.48 | 4.64 | 1.62E-05 |
| 4HG0337930 | F2DU32 | Predicted protein | -1.48 | 2.55 | 2.04E-02 |
| 6HG0559110 | A0A287TJQ3 | Uncharacterized protein | -1.50 | 4.57 | 1.40E-05 |
| 2HG0099060 | A0A8I6W949 | Chalcone synthase | -1.51 | 2.16 | 3.52E-02 |
| 6HG0614660 | A0A8I6YV57 | Pentacotripeptide-repeat region of PRORP domain-containing protein | -1.51 | 2.83 | 9.13E-03 |
| 2HG0206630 | A0A8I6WWY3 | Uncharacterized protein | -1.51 | 2.52 | 1.82E-02 |
| 2HG0178080 | A0A8I6XF77 | Uncharacterized protein | -1.52 | 4.67 | 7.99E-06 |
| 2HG0194770 | A0A8I6XH73 | Pectinesterase inhibitor domain-containing protein | -1.52 | 5.10 | 2.27E-06 |
| 7HG0752910 | A0A8I6YIK0 | Protein FAR1-RELATED SEQUENCE | -1.53 | 3.52 | 8.02E-04 |
| 7HG0728850 | F2CTI6 | Predicted protein | -1.55 | 3.13 | 2.79E-03 |
| 6HG0599280 | A0A8I6YKE5 | Aldehyde dehydrogenase | -1.57 | 2.26 | 2.11E-02 |
| 2HG0206030 | F2CZ63 | Predicted protein | -1.58 | 3.39 | 7.77E-04 |
| 7HG0635330 | A0A8I6Y013 | Uncharacterized protein | -1.59 | 2.31 | 1.81E-02 |
| 1HG0088680 | F2CXM2 | Uncharacterized protein ycf23 | -1.60 | 6.19 | 2.45E-07 |
| 1HG0087820 | F2D4R6 | Predicted protein | -1.60 | 4.59 | 2.77E-06 |
| 2HG0173380 | A0A8I6X4T9 | Amino acid transporter transmembrane domain-containing protein | -1.62 | 2.39 | 1.23E-02 |
| 2HG0122530 | A0A8I7B5A5 | Uncharacterized protein | -1.62 | 2.40 | 1.20E-02 |
| 3HG0301990 | M0XFT5 | Bifunctional inhibitor/plant lipid transfer protein/seed storage helical domain-containing protein | -1.63 | 6.05 | 1.29E-07 |
| 5HG0475370 | A0A8I6Y0V1 | isochorismate synthase (EC 5.4.4.2) | -1.63 | 2.30 | 1.46E-02 |
| 3HG0282170 | A0A8I6WSU8 | phosphoethanolamine N-methyltransferase (EC 2.1.1.103) | -1.63 | 7.61 | 2.27E-06 |
| 1HG0018050 | A0A8I6WN02 | Uncharacterized protein | -1.64 | 4.02 | 2.15E-05 |
| 1HG0012630 | F2CZJ1 | Predicted protein | -1.67 | 3.69 | 7.08E-05 |
| 6HG0545500 | M0YYX8 | Papain-like cysteine proteinase | -1.68 | 2.04 | 1.84E-02 |
| 6HG0542050 | A0A8I6Y952 | Beta-fructofuranosidase | -1.69 | 3.46 | 1.80E-04 |
| 7HG0702660 | A0A8I6YFS7 | 4-coumarate--CoA ligase (EC 6.2.1.12) | -1.69 | 2.43 | 7.26E-03 |
| 3HG0235910 | A0A8I6WX70 | FAS1 domain-containing protein | -1.83 | 4.94 | 8.25E-09 |
| 1HG0059470 | A0A8I6WTX7 | Uncharacterized protein | -1.84 | 6.35 | 2.04E-09 |
| 2HG0133180 | F2D4W9 | Ribosome-recycling factor, chloroplastic (Ribosome-releasing factor, chloroplastic) | -1.87 | 4.37 | 7.71E-08 |
| 3HG0241690 | M0UPD8 | Peroxidase (EC 1.11.1.7) | -1.92 | 2.14 | 3.64E-03 |
| 7HG0711020 | F2CWL8 | Predicted protein | -1.96 | 4.83 | 9.78E-10 |
| 3HG0218800 | F2DH05 | Predicted protein | -1.98 | 4.88 | 4.12E-10 |
| 1HG0000490 | F2CSQ0 | Chalcone-flavonone isomerase family protein | -2.02 | 3.51 | 2.86E-06 |
| 6HG0629290 | A0A8I6YGS3 | Alpha/beta hydrolase fold-3 domain-containing protein | -2.02 | 3.35 | 8.61E-06 |
| 7HG0640130 | A0A8I6YA44 | Flavonoid 3',5'-hydroxylase | -2.05 | 2.83 | 1.09E-04 |
| 4HG0357240 | F2DSS3 | Laccase (EC 1.10.3.2) (Benzenediol:oxygen oxidoreductase) (Diphenol oxidase) (Urishiol oxidase) | -2.08 | 2.97 | 3.69E-05 |
| 4HG0385330 | F2D677 | Glycosyltransferases (EC 2.4.-.-) | -2.08 | 2.38 | 5.38E-04 |
| 3HG0302520 | F2CPW9 | Laccase (EC 1.10.3.2) (Benzenediol:oxygen oxidoreductase) (Diphenol oxidase) (Urishiol oxidase) | -2.08 | 3.75 | 2.05E-07 |
| 5HG0509030 | M0ZCV6 | Pentatricopeptide repeat-containing protein | -2.09 | 2.37 | 5.38E-04 |
| 3HG0306130 | A0A8I6XA83 | Trigger factor ribosome-binding bacterial domain-containing protein | -2.12 | 2.87 | 4.21E-05 |
| 2HG0104780 | A0A287H227 | Ribulose bisphosphate carboxylase small subunit, chloroplastic (RuBisCO small subunit) | -2.15 | 7.07 | 1.47E-11 |
| 5HG0497820 | A0A8I6Y4S4 | Polysaccharide biosynthesis domain-containing protein | -2.20 | 2.54 | 1.06E-04 |
| 7HG0702510 | A0A8I6YHB9 | Uncharacterized protein | -2.20 | 2.96 | 1.23E-05 |
| 6HG0540930 | A0A8I7BAT6 | NmrA-like domain-containing protein | -2.21 | 2.96 | 1.11E-05 |
| 7HG0715300 | F2E9I7 | Predicted protein | -2.22 | 2.08 | 5.76E-04 |
| 2HG0096220 | F2D2G3 | Predicted protein | -2.25 | 2.51 | 8.17E-05 |
| 1HG0051100 | A0A8I6WG57 | AB hydrolase-1 domain-containing protein | -2.25 | 2.74 | 2.65E-05 |
| 3HG0222380 | F2DGF0 | Predicted protein | -2.26 | 4.35 | 6.02E-11 |
| 4HG0388460 | A0A8I6Y820 | Laccase (EC 1.10.3.2) (Benzenediol:oxygen oxidoreductase) (Diphenol oxidase) (Urishiol oxidase) | -2.32 | 5.43 | 1.65E-14 |
| 4HG0384780 | A0A8I7BCI4 | Trichome birefringence-like N-terminal domain-containing protein | -2.36 | 2.16 | 1.56E-04 |
| 6HG0542040 | A0A8I6YQ26 | Beta-fructofuranosidase | -2.36 | 2.08 | 2.16E-04 |
| 2HG0104810 | A0A287H239 | Ribulose bisphosphate carboxylase small subunit, chloroplastic (RuBisCO small subunit) | -2.37 | 6.89 | 3.42E-14 |
| 5HG0537360 | A0A8I7BG18 | DUF547 domain-containing protein | -2.38 | 2.67 | 1.14E-05 |
| 6HG0612680 | A0A8I6YUY7 | Protein ODORANT1 | -2.38 | 2.27 | 8.45E-05 |
| 5HG0525430 | A0A8I6YN10 | Trichome birefringence-like N-terminal domain-containing protein | -2.39 | 4.65 | 2.70E-13 |
| 3HG0303550 | A0A8I6X9U8 | DNA-3-methyladenine glycosylase I | -2.42 | 4.02 | 7.53E-11 |
| 7HG0726310 | F2DSR5 | Predicted protein | -2.46 | 3.74 | 6.74E-10 |
| 2HG0104820 | A0A287H239 | Ribulose bisphosphate carboxylase small subunit, chloroplastic (RuBisCO small subunit) | -2.50 | 3.24 | 5.26E-08 |
| 7HG0664720 | A0A8I6YKT5 | Phytocyanin domain-containing protein | -2.55 | 3.30 | 1.70E-08 |
| 5HG0513740 | F2D4V2 | Predicted protein | -2.55 | 2.19 | 3.44E-05 |
| 5HG0482430 | F2CRG2 | Predicted protein | -2.60 | 7.26 | 5.93E-16 |
| 2HG0104760 | A0A287H239 | Ribulose bisphosphate carboxylase small subunit, chloroplastic (RuBisCO small subunit) | -2.64 | 2.53 | 2.78E-06 |
| 7HG0667240 | F2E2E6 | Predicted protein | -2.65 | 2.23 | 1.40E-05 |
| 7HG0639980 | F2E6J2 | Sucrose synthase (EC 2.4.1.13) | -2.69 | 3.30 | 2.53E-09 |
| 1HG0076150 | A0A8I6WX52 | NmrA-like domain-containing protein | -2.71 | 5.48 | 2.49E-19 |
| 7HG0746540 | F2DRR5 | Dirigent protein | -2.73 | 2.20 | 9.32E-06 |
| 1HG0073550 | M0VC31 | Laccase (EC 1.10.3.2) (Benzenediol:oxygen oxidoreductase) (Diphenol oxidase) (Urishiol oxidase) | -2.77 | 2.77 | 1.35E-07 |
| 1HG0070730 | A0A8I7B1B7 | Glycosyltransferase (EC 2.4.1.-) | -2.79 | 2.32 | 2.77E-06 |
| 4HG0336940 | F2DL00 | Predicted protein | -2.96 | 2.41 | 3.48E-07 |
| 4HG0334970 | M0WME5 | Plasma membrane ATPase (EC 7.1.2.1) | -3.06 | 3.03 | 4.10E-10 |
| 3HG0286930 | F2EGN4 | Predicted protein | -3.06 | 5.96 | 4.04E-24 |
| 3HG0302570 | F2DXF2 | Laccase (EC 1.10.3.2) (Benzenediol:oxygen oxidoreductase) (Diphenol oxidase) (Urishiol oxidase) | -3.11 | 2.59 | 2.15E-08 |
| 3HG0252270 | A0A287KQB5 | Germin-like protein | -3.13 | 3.36 | 3.34E-12 |
| 1HG0083720 | F2D8V2 | Predicted protein | -3.16 | 4.18 | 4.21E-18 |
| 3HG0222410 | A0A8I6XA13 | Bowman-Birk serine protease inhibitors family domain-containing protein | -3.20 | 3.11 | 3.65E-11 |
| 6HG0629260 | A0A8I6YF88 | glutathione transferase (EC 2.5.1.18) | -3.39 | 7.22 | 2.14E-25 |
| 3HG0330120 | A5YTR4 | Flavonoid O-methyltransferase (Predicted protein) | -3.72 | 4.24 | 6.39E-24 |
| 5HG0529880 | A0A8I7BFV0 | Uncharacterized protein | -3.73 | 2.80 | 4.35E-12 |
| 1HG0073500 | F2D9J2 | Laccase (EC 1.10.3.2) (Benzenediol:oxygen oxidoreductase) (Diphenol oxidase) (Urishiol oxidase) | -4.01 | 4.23 | 4.52E-26 |

Table S4 Differentially expressed WRKY transcription factors. C, control; A, AMF addition; P, Puccinia addition; AP, AMF plus Puccinia.

| **Comparison** | **Name** | **Gene ID** | **logFC** |
| --- | --- | --- | --- |
| C Vs A | HvWRKY17 | HORVU.MOREX.r3.1HG0029360 | 1.704 |
|  | HvWRKY43 | HORVU.MOREX.r3.1HG0071500 | 1.646 |
|  | HvWRKY19 | HORVU.MOREX.r3.1HG0080920 | 1.392 |
|  | HvWRKY20 | HORVU.MOREX.r3.1HG0080940 | 1.727 |
|  | HvWRKY27/55/87 | HORVU.MOREX.r3.1HG0090460 | 1.732 |
|  | HvWRKY26 | HORVU.MOREX.r3.1HG0090520 | 1.888 |
|  | HvWRKY30 | HORVU.MOREX.r3.2HG0123480 | 1.659 |
|  | HvWRKY21 | HORVU.MOREX.r3.3HG0276810 | 2.106 |
|  | HvWRKY50 | HORVU.MOREX.r3.3HG0304380 | 1.051 |
|  | HvWRKY48 | HORVU.MOREX.r3.4HG0333220 | 1.528 |
|  | HvWRKY22 | HORVU.MOREX.r3.5HG0474120 | 1.926 |
|  | HvWRKY1/38 | HORVU.MOREX.r3.6HG0568570 | 2.042 |
|  | HvWRKY2 | HORVU.MOREX.r3.7HG0743270 | 2.858 |
|  | HvWRKY23 | HORVU.MOREX.r3.7HG0743280 | 2.730 |
| P Vs AP | HvWRKY3 | HORVU.MOREX.r3.5HG0484070 | -2.160 |
|  | HvWRKY52 | HORVU.MOREX.r3.5HG0484060 | -1.657 |
|  | HvWRKY105 | HORVU.MOREX.r3.2HG0111890 | -1.230 |
|  | HvWRKY99 | HORVU.MOREX.r3.4HG0378680 | -1.469 |
|  | HvWRKY31/105 | HORVU.MOREX.r3.1HG0070890 | -1.441 |
| A Vs AP | HvWRKY31 | HORVU.MOREX.r3.1HG0070890 | 2.288 |
|  | HvWRKY43 | HORVU.MOREX.r3.1HG0071500 | 2.588 |
|  | HvWRKY69 | HORVU.MOREX.r3.1HG0088760 | 2.838 |
|  | HvWRKY12 | HORVU.MOREX.r3.2HG0096750 | 2.185 |
|  | HvWRKY105 | HORVU.MOREX.r3.2HG0111890 | 2.118 |
|  | HvWRKY7/96/97/98 | HORVU.MOREX.r3.2HG0192860 | 1.297 |
|  | HvWRKY63 | HORVU.MOREX.r3.3HG0237200 | 4.356 |
|  | HvWRKY37 | HORVU.MOREX.r3.3HG0237360 | 3.810 |
|  | HvWRKY64 | HORVU.MOREX.r3.3HG0273500 | 2.889 |
|  | HvWRKY21 | HORVU.MOREX.r3.3HG0276810 | -1.540 |
|  | HvWRKY51 | HORVU.MOREX.r3.3HG0278140 | 1.381 |
|  | HvWRKY45 | HORVU.MOREX.r3.3HG0286660 | 2.978 |
|  | HvWRKY48 | HORVU.MOREX.r3.4HG0333220 | -1.410 |
|  | HvWRKY33 | HORVU.MOREX.r3.4HG0380330 | 1.456 |
|  | HvWRKY22 | HORVU.MOREX.r3.5HG0474120 | 1.452 |
|  | HvWRKY3 | HORVU.MOREX.r3.5HG0484070 | -1.128 |
|  | HvWRKY92 | HORVU.MOREX.r3.6HG0551140 | 2.084 |
|  | HvWRKY1/38 | HORVU.MOREX.r3.6HG0568570 | 1.009 |
|  | HvWRKY60 | HORVU.MOREX.r3.7HG0650260 | 1.129 |
|  | HvWRKY2 | HORVU.MOREX.r3.7HG0743270 | 1.086 |
| C Vs P | HvWRKY17 | HORVU.MOREX.r3.1HG0029360 | 1.349 |
|  | HvWRKY31 | HORVU.MOREX.r3.1HG0070890 | 5.725 |
|  | HvWRKY43 | HORVU.MOREX.r3.1HG0071500 | 4.644 |
|  | HvWRKY19 | HORVU.MOREX.r3.1HG0080920 | 2.493 |
|  | HvWRKY20 | HORVU.MOREX.r3.1HG0080940 | 2.892 |
|  | HvWRKY69 | HORVU.MOREX.r3.1HG0088760 | 3.730 |
|  | HvWRKY27/55/87 | HORVU.MOREX.r3.1HG0090460 | 1.179 |
|  | HvWRKY26 | HORVU.MOREX.r3.1HG0090520 | 2.456 |
|  | HvWRKY56 | HORVU.MOREX.r3.1HG0092420 | 3.134 |
|  | HvWRKY12 | HORVU.MOREX.r3.2HG0096750 | 3.142 |
|  | HvWRKY105 | HORVU.MOREX.r3.2HG0111890 | 3.934 |
|  | HvWRKY4/106 | HORVU.MOREX.r3.2HG0111920 | 1.456 |
|  | HvWRKY30 | HORVU.MOREX.r3.2HG0123480 | 2.616 |
|  | HvWRKY10 | HORVU.MOREX.r3.2HG0158690 | 1.664 |
|  | HvWRKY7/96/97/98 | HORVU.MOREX.r3.2HG0192860 | 1.884 |
|  | HvWRKY63 | HORVU.MOREX.r3.3HG0237200 | 4.887 |
|  | HvWRKY47 | HORVU.MOREX.r3.3HG0251900 | 1.583 |
|  | HvWRKY64 | HORVU.MOREX.r3.3HG0273500 | 3.205 |
|  | HvWRKY21 | HORVU.MOREX.r3.3HG0276810 | 1.158 |
|  | HvWRKY51 | HORVU.MOREX.r3.3HG0278140 | 2.401 |
|  | HvWRKY45 | HORVU.MOREX.r3.3HG0286660 | 4.467 |
|  | HvWRKY50 | HORVU.MOREX.r3.3HG0304380 | 1.880 |
|  | HvWRKY58 | HORVU.MOREX.r3.4HG0340900 | 1.049 |
|  | HvWRKY99 | HORVU.MOREX.r3.4HG0378680 | 2.449 |
|  | HvWRKY33 | HORVU.MOREX.r3.4HG0380330 | 1.449 |
|  | HvWRKY22 | HORVU.MOREX.r3.5HG0474120 | 3.534 |
|  | HvWRKY52 | HORVU.MOREX.r3.5HG0484060 | 3.322 |
|  | HvWRKY3 | HORVU.MOREX.r3.5HG0484070 | 1.081 |
|  | HvWRKY42 | HORVU.MOREX.r3.5HG0489860 | 1.312 |
|  | HvWRKY15 | HORVU.MOREX.r3.5HG0511790 | 2.030 |
|  | HvWRKY92 | HORVU.MOREX.r3.6HG0551140 | 3.262 |
|  | HvWRKY1/38 | HORVU.MOREX.r3.6HG0568570 | 2.951 |
|  | HvWRKY60 | HORVU.MOREX.r3.7HG0650260 | 2.214 |
|  | HvWRKY2 | HORVU.MOREX.r3.7HG0743270 | 4.261 |
|  | HvWRKY23 | HORVU.MOREX.r3.7HG0743280 | 3.459 |

WRKY information sourced from (Liu et al. 2005; Xu et al. 2006; Mukhtar et al. 2008; Berri et al. 2009; Chen et al. 2009; Bhattarai et al. 2010; Gao et al. 2011, 2016; van Verk, Bol and Linthorst 2011; Wei et al. 2013; Ding et al. 2014; Han et al. 2014; Huangfu et al. 2016; Jiang and Yu 2016; YANG et al. 2016; Yokotani et al. 2018; Dangol et al. 2021; Huang et al. 2021; Wang et al. 2021, 2023, 2024; Li et al. 2024; Son, Song and Im 2024; Zhu et al. 2025)

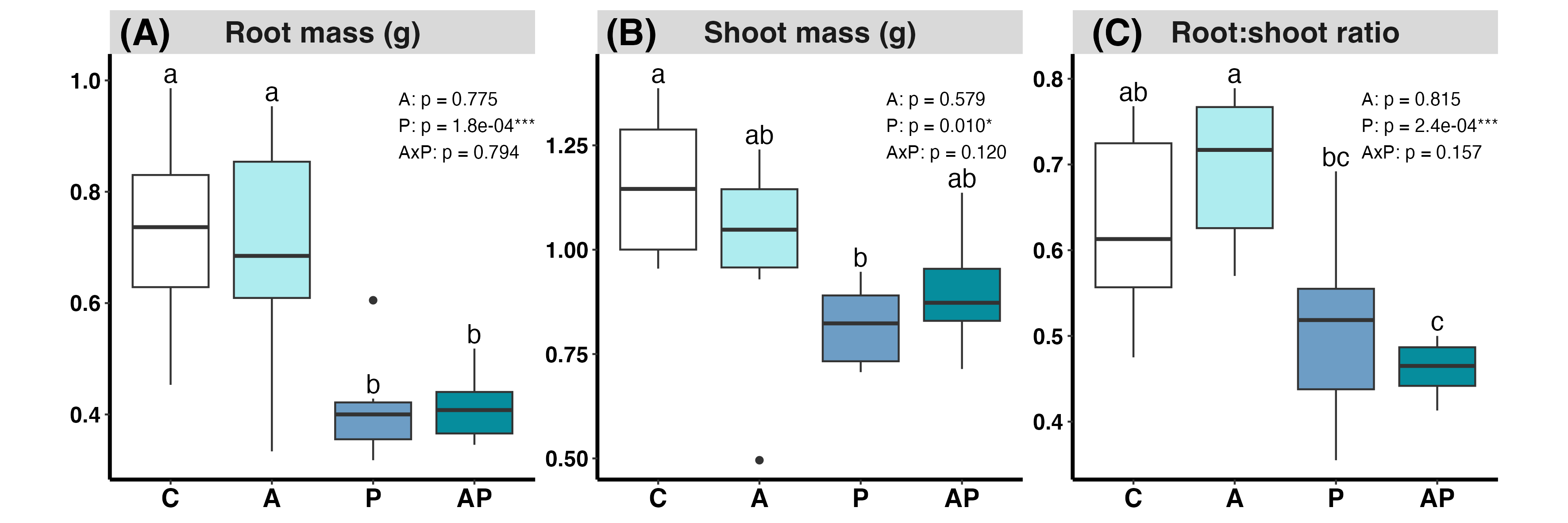

Figure S1 **Effects of AMF inoculation and** Puccinia **infection on barley biomass allocation.**
Boxplots show **(A)** root dry mass (g), **(B)** shoot dry mass (g), and **(C)** root:shoot ratio of barley plants grown under four treatments: Control (C), AMF inoculated (A), Puccinia-infected (P), and combined AMF inoculation and Puccinia infection (AP). Boxes represent the interquartile range with the median indicated by the horizontal line; whiskers extend to 1.5 × the interquartile range, with individual points representing outliers. Different letters above boxes indicate significant differences among treatment combinations based on post-hoc comparisons of estimated marginal means. Effects of AMF inoculation (A), Puccinia infection (P), and their interaction (A×P) were tested using two-way ANOVA; corresponding p-values (with significance indicated by asterisks: * p < 0.05, ** p < 0.01, *** p < 0.001) are shown within each panel (n = 6).

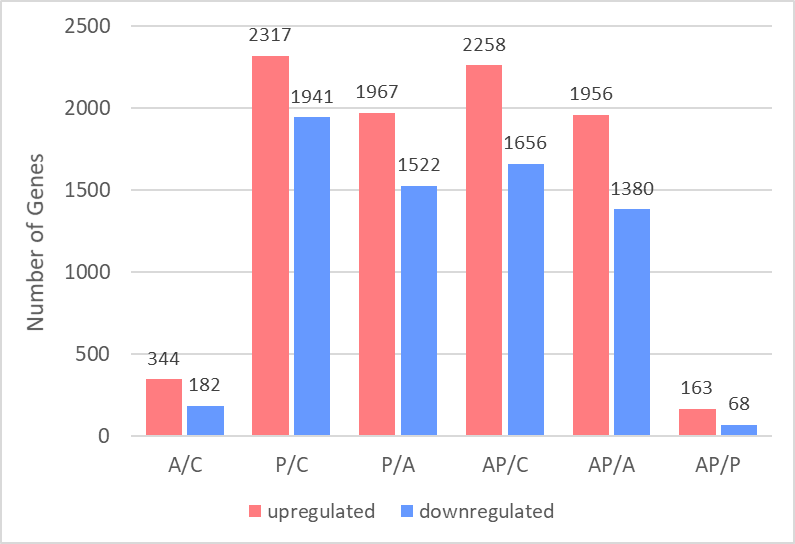

Figure S2 **Differentially expressed genes across treatment contrasts.**
Bars show the number of significantly upregulated (red) and downregulated (blue) genes identified by RNA-seq for pairwise contrasts between treatments: Control (C), AM fungi (A), Puccinia hordei (P), and AM fungi plus P. hordei (AP). Differential expression was determined using edgeR with genes defined as differentially expressed at a false discovery rate (FDR) < 0.05, fold change > 1, and logCPM > 2. Numbers above bars indicate the total number of genes in each category for the corresponding contrast.

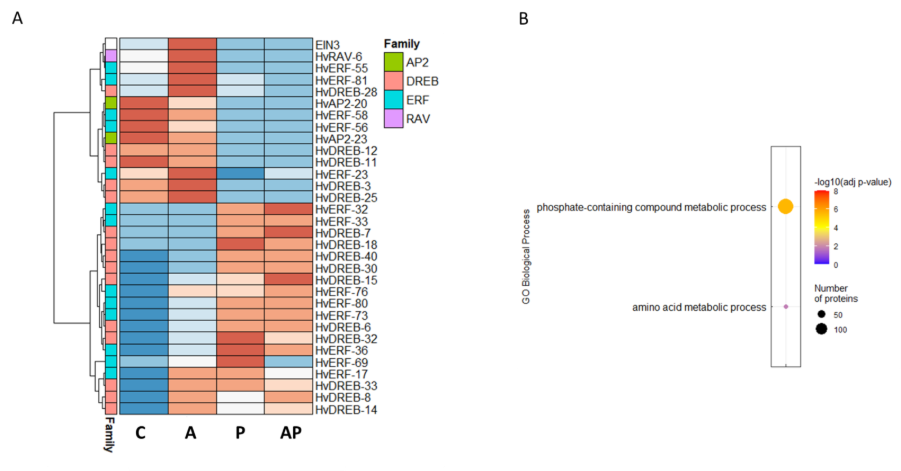

Figure S3 Expression and GO analysis of ethylene signalling genes. **(A)** Scaled expression of differentially expressed ethylene signalling genes: Ethylene Insensitive 3 (EIN3), Ethylene Response Factors (ERFs), Dehydration-Responsive Element Binding (DREB), Related to ABI3/VP1 (RAV), and APETALA2 (AP2) in barley leaves under four treatments: control – C, AMF – A, Puccinia – P, and AMF + Puccinia – AP. (FC > 2, FDR < 0.05, logCPM > 2). **(B)** GO enrichment of 1,321 DEGs with ERF-binding motifs in promoter regions (FIMO, p < 0.0001).

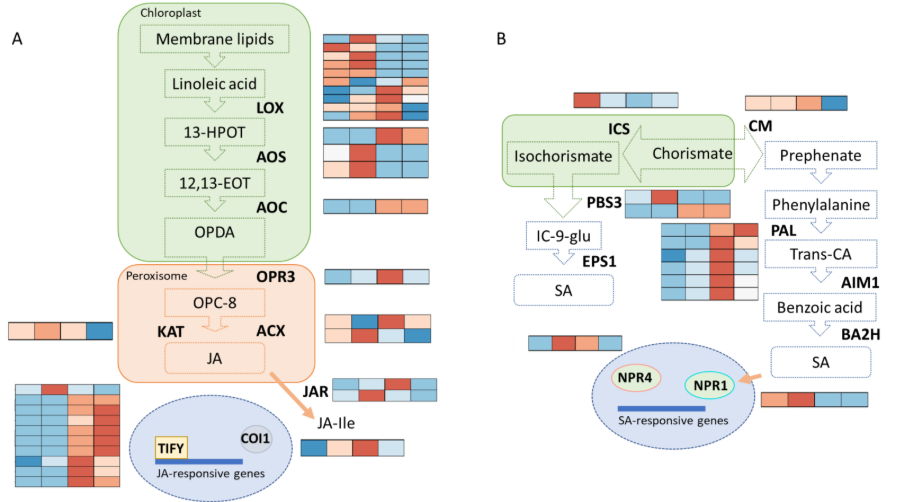

Figure S4 J**asmonic acid (JA) and salicylic acid (SA) signalling gene expression.** (A) Heatmap of jasmonic acid biosynthesis and signalling genes in barley leaves under control (C), AMF (A), Puccinia (P), and AMF + Puccinia (AP) treatments. Genes include Lipoxygenase (LOX), Allene Oxide Synthase (AOS), Allene Oxide Cyclase (AOC), Oxophytodienoate Reductase 3 (OPR3), Ketoacyl-CoA Thiolase (KAT), Acyl-CoA Oxidase (ACX), Jasmonate Resistant (JAR), and Coronate Insensitive 1 (COI1). (B) Heatmap of salicylic acid biosynthesis and signalling genes across the same treatments. Genes include Isochorismate Synthase (ICS), Chorismate Mutase (CM), Avrpphb Susceptible 3 (PBS3), Enhanced Pseudomonas Susceptibility 1 (EPS1), Phenylalanine Ammonia Lyase (PAL), Abnormal Inflorescence Meristem 1 (AIM1), Benzoic Acid 2-Hydroxylase (BA2H), and Non-Expressor of Pathogenesis-Related 1 (NPR1).

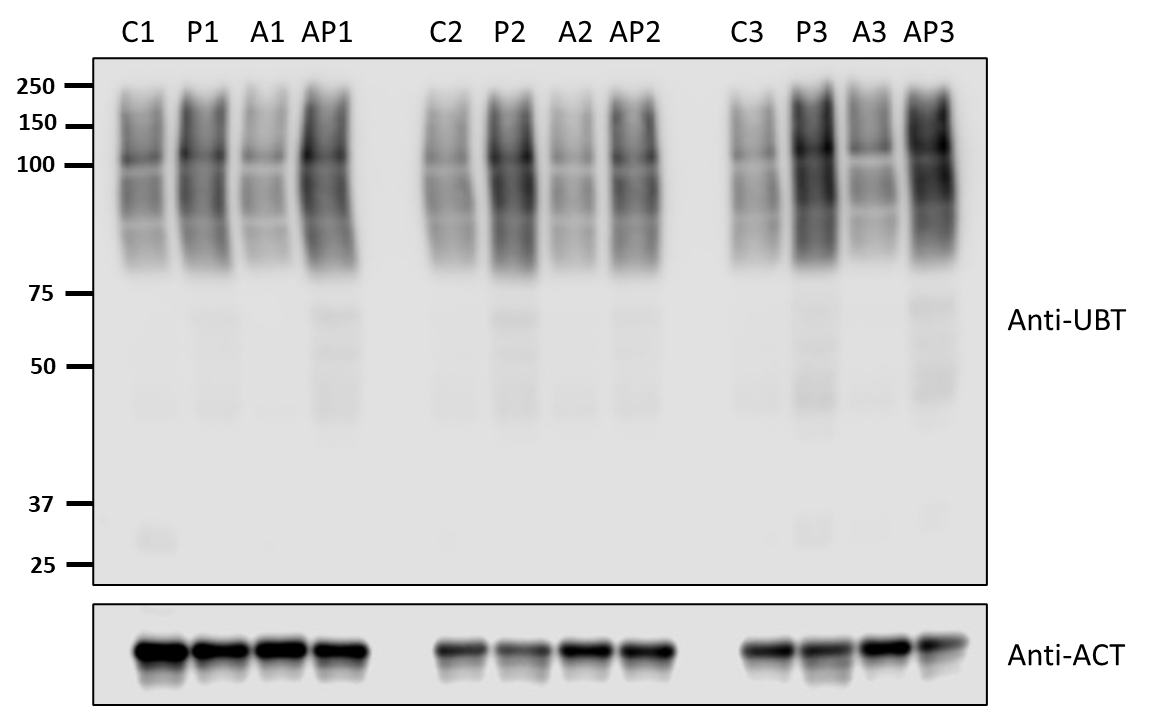

Figure S5 Western blot of the ubiquitin signal normalised by actin in Control, Puccinia-infected, AMF-colonised and AMF-colonised Puccinia-infected plants showing all replicates. C, control; A, AMF addition; P, Puccinia addition; AP, AMF plus Puccinia (n = 3).

1. ^*^ These authors contributed equally to this work
2. ^†^ Corresponding author: Thorunn Helgason,

   Institute of Ecology & Evolution,

   School of Biological Sciences,

   University of Edinburgh,

   Edinburgh, EH9 3BF, UK
